# Regulation of Small RNAs by Exercise and Their Role in Insulin Sensitivity

**DOI:** 10.64898/2026.05.12.724616

**Authors:** Christopher G. Vann, Akshay Bareja, Monica J. Hubal, Syeda Iffat Naz, Sisi Ma, Melissa C. Orenduff, Leanna M. Ross, William C. Bennett, Kim M. Huffman, Constantin F Aliferis, William E. Kraus, Virginia B. Kraus

**Author notes:** Address correspondence to: Christopher G. Vann, PhD Research Scholar, Duke Molecular Physiology Institute, Virginia Byers Kraus, MD, PhD, Mary Bernheim Distinguished Professor of Medicine Professor of Orthopaedics and Pathology, Division of Rheumatology, Duke University School of Medicine Duke Molecular Physiology Institute.

## Abstract

We investigated effects of three aerobic exercise interventions, varying in amount and intensity with durations of 8–9-months on small RNA (smRNA) expression and regulatory pathways in skeletal muscle and plasma from 120 participants. Using untargeted smRNA sequencing focused on miRNAs and piRNAs, adjusting for demographics and bodyweight, we identified 124 muscle smRNAs altered by exercise amount and 15 by intensity, and 47 plasma smRNAs altered by intensity and one by amount. These smRNAs were enriched in metabolic, transcriptional, translational, and cell cycle pathways. Exercise-induced changes in several smRNAs–six from muscle and five from plasma–and exercise-induced reduction in body weight, aligned with improvement in insulin sensitivity (p<0.05). These findings demonstrate tissue-specific regulation of smRNAs by exercise and identify potential candidates for exercise mimetics to modulate muscle insulin sensitivity.

## INTRODUCTION

Exercise improves health and performance, partly through adaptations in skeletal muscle, the largest organ system in the human body. This system is primarily responsible for postural control, locomotion, and thermogenesis, while also playing critical roles in whole-body glucose metabolism and energy homeostasis^1,2^. Exercise regulates RNA expression, including small non-coding RNAs (smRNAs), across multiple organ systems^3^.

Whole-body glucose metabolism is regulated, in part, by the coordinated actions of insulin and glucagon^4^. Regular exercise reduces both fasting and glucose-stimulated insulin levels^5,6^. In the first Studies of Targeted Risk Reduction Intervention through Defined Exercise (STRRIDE I) randomized trial^7^, 8-9 months of supervised aerobic exercise improved skeletal muscle (SkM) insulin sensitivity, assessed by intravenous glucose tolerance testing (IVGTT)^8^ – an effect reflected in reduced circulating insulin concentrations in response to a dynamic metabolic challenge^9^. Given that insulin signaling is regulated by some microRNAs (miRNAs)^10^, their identification through an untargeted approach, as performed here, may reveal specific miRNAs that modulate insulin action and mediate the metabolic benefits of exercise.

Regulatory smRNAs are a major class of epigenetic modifiers that generally act to suppress gene expression through translational repression or degradation of target messenger RNAs (mRNAs)^11–13^. This class includes miRNAs (15-25 nt length) and piwi-interacting RNAs (piRNAs; 22-35 nt). MiRNAs regulate approximately 50-60% of protein-coding genes, while piRNAs maintain germline DNA and silence transposable elements^14–17^. Analogous to miRNAs, piRNAs may silence gene expression through their interaction with PIWI proteins to form silencing complexes^18,19^. Since their discovery in the early 2000s^20^, piRNAs have remained incompletely characterized; however, recent advances in sequencing technologies and computational methods are revealing a broader spectrum of functions for this class of smRNAs^21^, including their potential utility as circulating plasma biomarkers^22^.

Many studies, both *in vivo*^23–25^ and *in vitro*^26^, have identified exercise responsive smRNAs playing roles in muscle physiology and health. Several studies have investigated the dose-response relationship of physical activity on various health outcomes by manipulating frequency, amount, or intensity, but evidence from clinical trials remains sparse^27^. To our knowledge, the independent effects of exercise amount and intensity on SkM and plasma smRNA expression have not been previously evaluated. We aimed to address this knowledge gap through analyses of the STRRIDE I and II trials.

Designed to understand the individual and combined effects of aerobic exercise, amounts and intensities, on cardiometabolic risk factors, the STRRIDE trials demonstrated dose-responsive benefits of aerobic exercise on key metabolic health markers, including reductions in subcutaneous and visceral adiposity, improved blood lipid profiles and insulin sensitivity, and mitigation of metabolic syndrome features^28–33^. Specifically, a greater amount of exercise results in improvements in HDL cholesterol^34^ and reductions in abdominal adiposity^31^, whereas exercise intensity is linked to improvements in fasting insulin^35^ and circulating triglycerides^34^. Notably, when energy expenditure was matched, moderate-intensity exercise produces more robust improvements in insulin sensitivity than vigorous-intensity exercise^35^. Thus, the design of the STRRIDE trials offers a unique opportunity to investigate the role of smRNA expression in the context of these dose-responsive, exercise-induced health effects. Leveraging data and samples from STRRIDE I and II participants, randomized to either the inactive control group or one of three aerobic exercise groups, our goal was to identify smRNAs modified by exercise and, in turn, the biological pathways they regulate, as well as their relationship to a key clinical outcome, change in insulin sensitivity. We hypothesize that exercise amount and intensity will have differing effects on smRNA expression patterns and that exercise-induced metabolic adaptations are mediated, at least in part, through small (sm)RNA-associated inter-organ signaling.

## MATERIALS AND METHODS

### Ethical Approval

All protocols and procedures for the STRRIDE I (NCT00200993; 1999-2003)^7^ and STRRIDE II (“STRRIDE AT/RT”; NCT00275145; 2004-2008)^36^ clinical trials were approved by institutional review boards at Duke University and East Carolina University. This study conformed to the standards set by the latest revision of the declaration of Helsinki. All study participants provided oral and written informed consent.

### STRRIDE I and II Study Designs

STRRIDE I and STRRIDE II (AT/RT) enrolled previously sedentary men and women aged 18-70 years with overweight or class I obesity (BMI 25-35 kg/m^2^) and mild-to-moderate dyslipidemia (low-density lipoprotein-cholesterol (LDL-C) 130-190 mg/dL or high-density lipoprotein cholesterol (HDL-C) ≤40 mg/dL for men and ≤45 mg/dL for women). Upon study enrollment, participants were instructed to maintain current lifestyle factors, including dietary intake and medication use, and refrain from exercise outside of the intervention. Participants were randomly assigned to either a non-exercise control group (CTL) for 6-months or a supervised exercise intervention group for 8-9-months, with biological samples collected at multiple timepoints throughout the intervention^7,28^. For the current investigation, we focused on participants who completed CTL or one of the aerobic exercise interventions in STRRIDE I and II, with matched SkM and plasma samples collected before (PRE) and after (POST) the study, resulting in a total of 120 participants (total sample mean age of 51.3±7.6; 47.5% female) assigned to one of the following groups:

1. CTL (n=11)
2. Low amount, moderate intensity aerobic exercise (LowMod; n=16): 14 kcal of energy expenditure/kg of body weight/week (KKW) at 40-55% peak oxygen consumption (̇VO_2 peak_)
3. Low amount, vigorous intensity aerobic exercise (LowVig; n=64): 14 KKW at 65-80% ̇VO_2peak_
4. High amount, vigorous intensity aerobic exercise (HighVig; n=29): 23 KKW at 65-80% ̇VO_2peak_.

For the aerobic training groups, exercise intensity and duration for all exercise sessions were verified by direct supervision and/or with the use of downloadable heart rate monitors (Polar Electro, Woodbury, NY, United States). Training adherence was calculated for each participant as the number of minutes completed within the prescribed heart rate range, divided by the number of total weekly minutes prescribed. The adherence percentage recorded for each attended exercise session was multiplied by the weekly caloric expenditure prescription to determine the achieved exercise amount. The achieved exercise amounts ranged from 533.0 to 2332.0 kCal/week (mdn=1231.3; Q_1_=1092; Q_3_=1382.2) expended. For analysis, exercise intensity was coded as 0 (inactive), 1 (moderate intensity), or 2 (vigorous intensity), corresponding to no exercise (CTL), or exercise performed at approximately 50% ̇VO_2peak_, or 75% ̇VO_2peak_, respectively. Importantly, heart rate and oxygen consumption during submaximal exercise show a strong linear relationship, making heart rate an appropriate proxy for ̇VO_2_ during aerobic exercise trials and a practical measure for monitoring participant compliance with the prescribed exercise intensity. Detailed participant characteristics are provided in **Table 1**, with study design illustrated in **Figure 1**.

**Figure 1:**
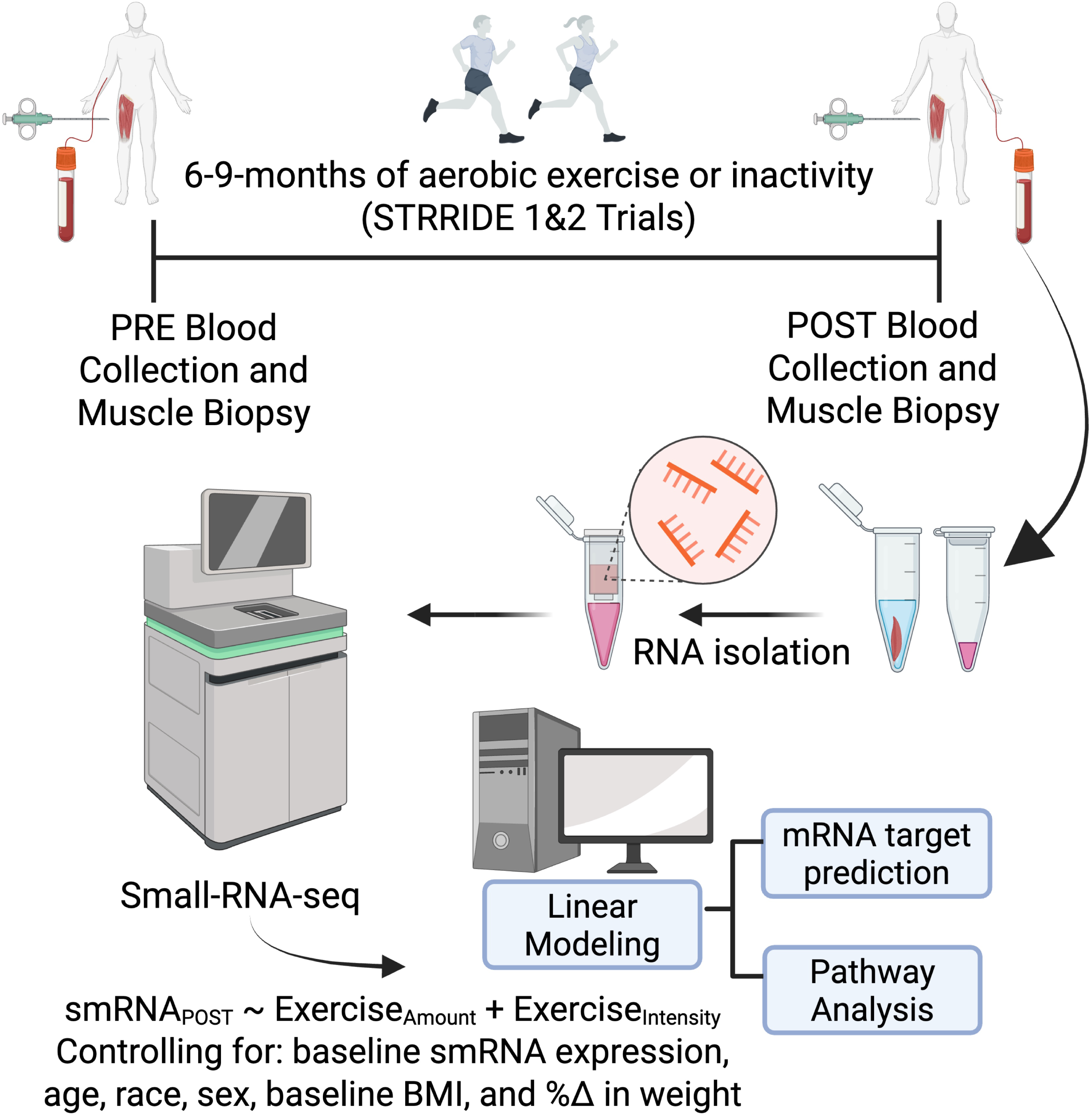
Experimental Design—a Global Overview of the STRRIDE Intervention and Biospecimens Used. Abbreviations: Small-RNA-seq, small RNA sequencing. This figure represents a global overview of the STRRIDE intervention and biospecimens used. Participants performed 6-9 months of inactivity or aerobic exercise of various amounts and intensities. Linear regression modeling controlled for baseline smRNA expression, age, race, biological sex, baseline body mass index (BMI_PRE_), and percent change in bodyweight. The experimental design figure was created using BioRender.com

**Table 1:** Participant Demographics.

| Variable | STRIDE I (n=71) | STRIDE II (n=49) | Total (n=120) |
| --- | --- | --- | --- |
| **Sex, n= (%)** |  |  |  |
| Male | 39 (54.9) | 24 (49.0) | 63 (52.5) |
| Female | 32 (45.1) | 25 (51.0) | 57 (47.5) |
| **Race, n= (%)** |  |  |  |
| White | 53 (74.6) | 41 (83.7) | 94 (78.3) |
| Black | 15 (21.1) | 8 (16.3) | 23 (19.2) |
| Other | 3 (4.2) | 0 (0.0) | 3 (2.5) |
| **Age, years (Range)** | 51.9 (40 - 66) | 50.5 (25 - 68) | 51.3 (25 - 68) |
| **BMI<sub>PRE</sub>, kg/m<sup>2</sup> (Range)** | 29.23 (24.4 - 38.1) | 30.5 (25.1 - 37.5) | 29.8 (24.4 - 38.1) |
| **%Δ BW, kg (Range)** | -0.81 (-10.26 - 7.70) | -1.70 (-11.91 - 7.41) | -1.17 (-11.91 - 7.70) |
| Exercise Variables |  |  |  |
| **Intensity (n=)** |  |  |  |
| Inactive (0) | 11 | 0 | 11 |
| Moderate (1) | 16 | 0 | 16 |
| Vigorous (2) | 44 | 49 | 93 |
| **BioAmount (Range)** | 1403.1 (533.0 - 2332.0) | 1329.3 (910.0 - 2173.6) | 1369.9 (533.0 - 2332.0) |

BMI, body mass index; BW, bodyweight; CTL, control. Data are presented as sample size with percentage of cohort representing each variable for sex and race. Intensity is presented as sample size for each intensity range. BioAmount is the product of assigned exercise amount (kCal) and intervention adherence percentage. Age, BMI<sub>PRE</sub>, percent change in BW, and BioAmount are presented as mean plus lower and upper bound range of the data.

### Laboratory Measurements and Collection of Biological Samples

#### Anthropometrics

With the participant wearing lightweight clothing and no shoes, height and body weight were measured with a stadiometer and digital scale, respectively.

#### Insulin Sensitivity

IS_i_ was determined using a three-hour IVGTT^8^. Through an intravenous catheter placed in the antecubital space, glucose (50% at 0.3 g/kg body mass) was injected at time zero and insulin (0.025 U/kg body mass) was injected at *minute 20.* Twenty-nine blood samples (at *minutes 0, 2, 3, 4, 5, 6, 8, 10, 12, 14, 16, 19, 22, 23, 24, 25, 27, 30, 40, 50, 60, 70, 80, 90, 100, 120, 140, 160, 180*) were obtained, centrifuged, and stored at -80 °C. Insulin was measured by immunoassay (Access Immunoassay System, Beckman Coulter, Fullerton, CA), and glucose with an oxidation reaction (YSI 2300, Yellow Springs, OH). IS_i_ was calculated using Bergman’s minimal model^8^. The IVGTT was performed after an overnight fast both at baseline and POST (16–24 h after the last training session for exercisers).

#### SkM Biopsies

SkM biopsies were taken from the vastus lateralis muscle via standard Bergstrom needle biopsy; approximately 100-200 mg of tissue was obtained with a triple pass of the needle and flash frozen^37^. SkM biopsies were performed after an overnight fast both at baseline and POST (16–24 h after the last training session for exercisers).

#### SkM and Plasma RNA Extraction and small RNA Sequencing

We extracted total RNA from ∼10 mg of SkM for each timepoint using a commercially available kit (Qiagen AllPrep DNA/RNA/miRNA extraction kits (Cat ID: 80224; Germantown, MD, USA) in accordance with the manufacturer’s specifications with slight modification during the incubation phases; all kits and reagents were from the same lot. Specifically, per sample, a cocktail consisting of 1 μL of RNA grade glycogen (Thermo Fisher; Waltham, MA, Cat ID: R0551) was added to 70 μL Buffer RDD and 10 μL DNAse I with the cocktail being added to each column and incubated at room temperature for 15 minutes prior to finishing the isolation as directed. Resulting RNA samples were quantified using the Qubit High Sensitivity RNA Quantification assay (Thermo Fisher, Cat ID: Q32855) and stored at -80°C until further processing. Total RNA from PRE and POST fasting plasma samples were isolated using the Qiagen miRNeasy Serum/Plasma Advanced Isolation Kit (Cat ID: 217204) in accordance with the manufacturer’s specifications; all kits and reagents were from the same lot. Resultant RNA samples were quantified using the Qubit High Sensitivity RNA Quantification Assay and then stored at - 80°C until further processing.

Total RNA from SkM (360 ng at 20 ng/μL) and plasma (15 μL) samples was submitted to the Duke Center for Genomic and Computational Biology for smRNA-sequencing (smRNAseq). Libraries were prepared using QIAseq miRNA Library Kits (Qiagen; Cat ID: 331502) in accordance with the manufacturer’s specifications. Sequencing was completed at 100bp, single end at a depth of 30M via Illumina NovaSeq6000. Importantly, all samples were submitted as a single submission with library prep kits and flow cells coming from the same lot.

### Data Processing and Analysis

FASTQ files generated via smRNA sequencing were mapped using the Qiagen GeneGlobe miRNA Primary Quantification Server using the Legacy 2.0 pipeline. This pipeline aligns raw reads to the Hg38 version of the human genome and collapses nested sequences into a single readout. CutAdapt was used to trim the 3’ adapter and remove low quality bases. The unique molecular identifiers (UMI)-tools package was used to identify insert sequences and UMIs; reads with <16bp insert sequences (i.e., reads that are too short) and UMI sequences <10bp (defective UMI) were discarded. MiRNA sequences were mapped to miRbase (v21) while piRNA sequences were mapped to piRNABank using bowtie2. FeatureCounts was used to quantify raw reads, which were exported and read into R (v4.2) for further processing.

Following the import of raw counts, data were log_2_CPM-transformed to account for differences in library size and filtered to remove low expressors (row sum <50 counts for a given smRNA). Explained variance of the exercise modified smRNA was assessed using variancePartition^38^ by fitting a linear model for each smRNA, with log_2_CPM expression as the outcome variable and Exercise_AMOUNT_, Exercise_INTENSITY_, sex, race, age, BMI_PRE_, and percent change in bodyweight included as predictors. This approach enabled estimation of the proportion of variance in smRNA expression attributable to each predictor, while simultaneously adjusting for the others.

#### Statistical and bioinformatic analyses

We investigated the effects of exercise amount and intensity on smRNA expression profiles via linear regression modeling, using post-exercise smRNA expression (smRNA_POST_) as the outcome, with predictors of Exercise_AMOUNT_ and Exercise_INTENSITY_, controlling for baseline smRNA expression (smRNA_PRE_), age, race, sex, baseline BMI (BMI_PRE_), and percent change in bodyweight. This model allowed for the control of baseline smRNA expression, addressing the variability among participants at baseline as recommended by previous methodological guidance^39,40^. Because smRNA research in the context of exercise interventions is relatively new and few studies have applied sequencing-based approaches, we set significance for our primary linear modeling at p<0.05, with a Benjamini-Hochberg false discovery rate threshold of Q<0.1, balancing sensitivity with an acceptable false positive rate^41^. Pathway analysis was performed on all miRNAs from Exercise_AMOUNT_ and Exercise_INTENSITY_ analyses using the miRNA Enrichment Analysis and Annotation Tool (miEAA v2.1), employing the GSEA analysis module^42,43^. Prior to analysis, miRNAs were rank ordered based on their β-estimate; we required a minimum of two miRNAs to be present per identified pathway with significance set at p<0.05. Modeling of miRNA-to-mRNA regulatory networks was completed using miRNet v2^44^, leveraging the miRTarBase v9 database. Regulatory networks were filtered using two methods: (i) a minimum network filter, which retained seed nodes and necessary non-seed nodes to preserve connectivity, and (ii) a shortest path filter, which reduced the network to nodes and edges that lie on the shortest path to nodes of interest. PiRNA associations with genomic repeat elements were investigated using RepeatMasker; potential gene targets were identified via sequence complementarity using piRNAQuest v2.0, piRBase, and NCBI blastn using the reverse complement sequence, using the entire mature piRNA sequence, allowing for a single nucleotide mismatch for each piRNA-mRNA match^45–48^. Functional assessment of piRNA was completed with DAVID^49^ using predicted mRNA targets as the input. All piRNAs were identified and discussed by their NCBI accession ID (DQ ID), as this identifier is consistent across databases.

To investigate the relationship between plasma and SkM smRNA expression, we performed Spearman correlations on log_2_CPM fold change scores (log_2_FC; calculated as: log2 [^𝑃𝑂𝑆𝑇^/_𝑃𝑅𝐸_]) for each smRNA; the correlation was considered significant if p<0.05. To identify SkM and plasma smRNAs associated with change in ISi, we used univariable linear regression modeling with each variable being tested independently. For modeling, we used log_2_FC in IS_i_ as the outcome variable with log_2_FC smRNA (calculated as: log_2_ [^𝑃𝑂𝑆𝑇^/_𝑃𝑅𝐸_]); we also evaluated independent demographic variables (age, race, sex, percent change in bodyweight, and BMI_PRE_). Standardized β-estimates were derived by multiplying the unstandardized regression coefficient by the ratio of the standard deviation of the predictor to that of the outcome (Log_2_FC in IS_i_); variables with p<0.05 were considered significant. Figures were created using GraphPad Prism (v10.1.1) and ggplot2^50^; schematics were created using biorender.com.

## RESULTS

### Effects of Aerobic Exercise on SkM smRNA Expression

We used multivariable linear regression modeling to evaluate the effects of exercise amount and intensity on SkM smRNA expression controlling for baseline smRNA expression, age, race, sex, baseline body mass index (BMI), and percent change in bodyweight. We assessed change in SkM smRNA expression with exercise amount —accounting for intensity, while also assessing exercise intensity —accounting for amount (**Figure 2a; Supplemental Table S1**). Based on significance p<0.05, we identified 235 ‘amount’-responsive smRNAs (208 miRNAs, 27 piRNAs), of which 124 (114 miRNAs, 10 piRNAs) remained significant following Benjamini-Hochberg correction (Q<0.1; 117 upregulated, 7 downregulated). A total of 190 smRNAs were intensity-responsive (p<0.05; 118 miRNAs, 72 piRNAs), of which 15 (12 miRNAs, 3 piRNAs) remained significant based on a Q<0.1 significance threshold (3 upregulated, 12 downregulated).

**Figure 2:**
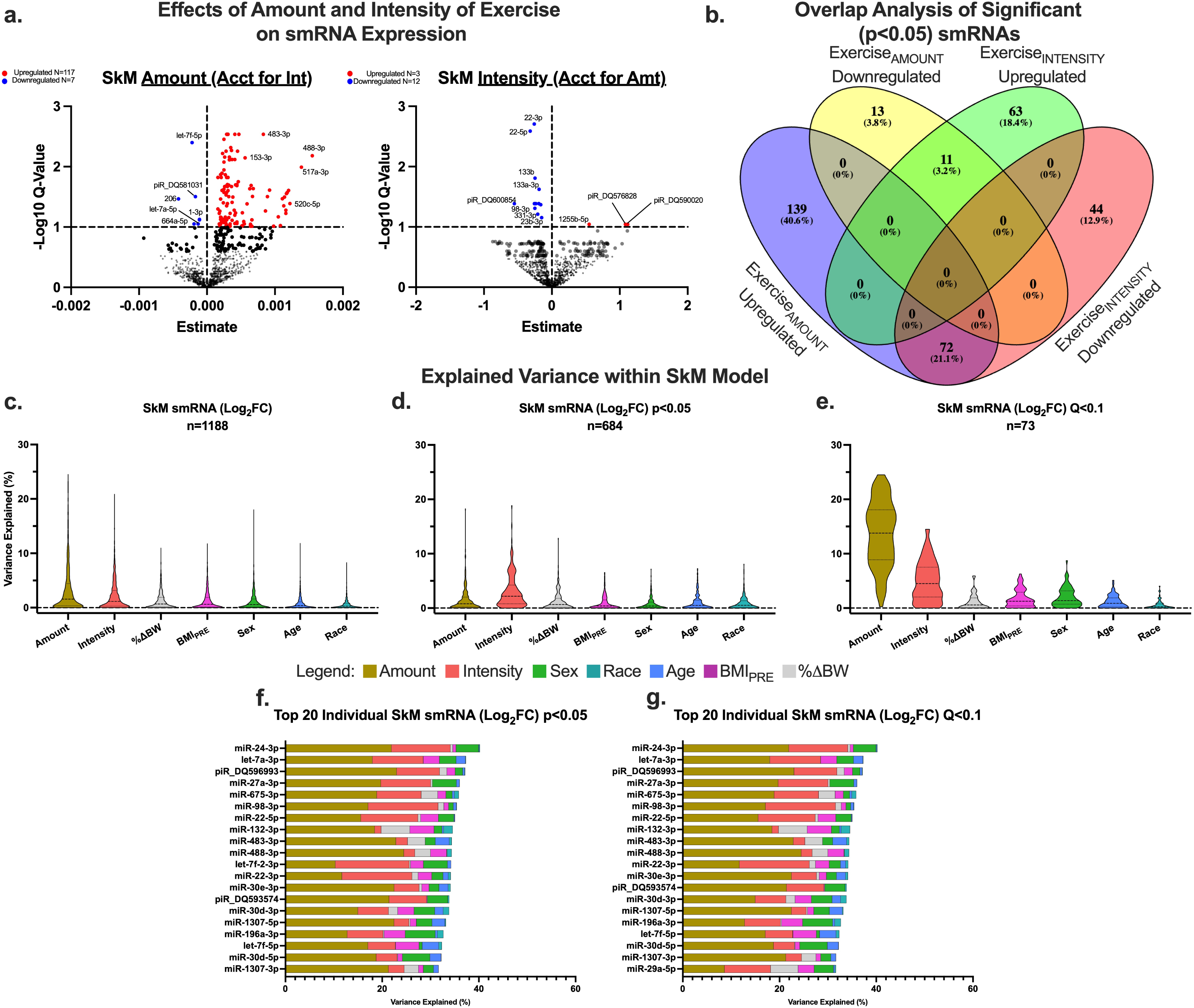
Effects of Exercise Amount and Intensity on smRNA Expression in Skeletal Muscle. Sample size: 120 (63 male, 57 female); Abbreviations: SkM, skeletal muscle; smRNA, small RNA; Amt, amount of exercise; Int, intensity of exercise; BMI_PRE_, Baseline BMI; %ΔBW, percent change in bodyweight. **Panel A** represents volcano plots of the -Log10 Q-value and estimate for adjusted SkM smRNA models. Non-significant SkM smRNAs are depicted in black, significantly upregulated (p<0.05; Q<0.1) SkM smRNAs are depicted in red, and significantly (p<0.05; Q<0.1) downregulated SkM smRNAs are depicted in blue. **Panel B** represents analysis of overlap whereas the Venn diagram shows the total number of nominally significant (p<0.05) up- and down-regulated SkM smRNAs for amount of exercise (accounting for intensity) and for intensity of exercise (accounting for amount) and the specific smRNAs that overlap in these variables. 72 smRNAs were identified as upregulated with exercise amount but downregulated with exercise intensity; 11 smRNAs were identified as downregulated by exercise amount but upregulated with exercise intensity. **Panel C-E** represent explainable variance with our linear modeling where violin plots depict analysis on all SkM smRNAs as well as smRNAs with p<0.05 and Q<0.1. **Panel F-G** are bar plots depicting variance in the top 20 SkM smRNAs with p<0.05 and Q<0.1; the remaining variance not depicted is attributable to unmeasured factors accounted for in residuals.

In addition, we identified 83 exercise-responsive smRNAs (p<0.05) independently associated with both exercise amount *and* intensity. In all cases, the effects of amount and intensity were divergent: 72 smRNAs were downregulated by intensity but upregulated by amount, whereas 11 showed the opposite pattern, being upregulated by intensity but downregulated by amount (**Figure 2b**). Interestingly, the majority of DE smRNAs upregulated by amount but downregulated by intensity were miRNAs (n=69). SmRNAs with the reverse pattern were predominantly piRNAs (n=8).

Using VariancePartition, our model explained ∼11% of the total variance in skeletal muscle smRNAs (n=1188; **Figure 2c; Supplemental Table S2**). Further, our model explained a mean 16.3% of the variance in the significant skeletal muscle exercise-responsive smRNAs (p<0.05; **Figure 2d**), with exercise amount accounting for 6.8% (range 0.0% to 24.5%) and exercise intensity 3.6% (range 0.0% to 20.9%). Among the most exercise-responsive smRNAs (passing the Q<0.1 threshold; **Figure 2e**), the explained variance increased to a mean 25.3%, with exercise amount accounting for 13.6% (range 0.0% to 24.5%) and intensity for 5.0% (range 0.0% to 14.5%). Notably, in top exercise-responsive SkM smRNAs (**Figure 2f-g**), exercise amount explained more than twice the variance of intensity, underscoring amount as the predominant factor influencing changes in SkM smRNA expression.

### Pathways Associated with Skeletal Muscle smRNAs

Pathway analysis of skeletal muscle miRNAs revealed over 1,300 enriched or depleted pathways (p <0.05), including metabolic, transcriptional, translational, and cell cycle processes, with both exercise amount and intensity yielding distinct signatures (**Figure 3a–b; Supplemental Table S3**). Notably, oxidative phosphorylation, a key metabolic pathway modulated by aerobic exercise involving 9 miRNAs targeting 31 mRNAs, was predicted to be depleted with increasing exercise amount (p=0.001) and enriched with greater exercise intensity (p=0.028) (**Figure 3c**).

**Figure 3:**
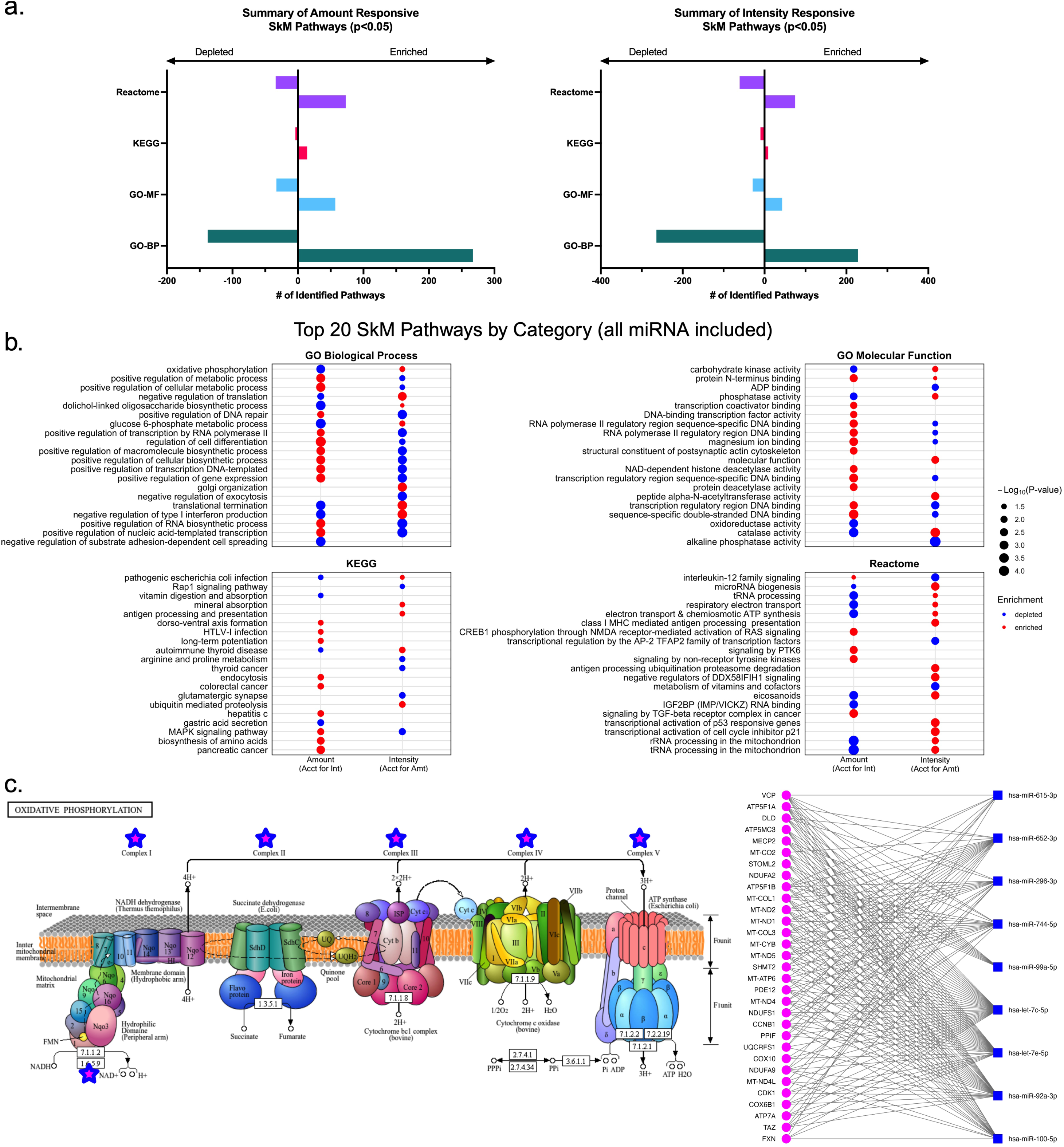
Pathways Associated with Skeletal Muscle smRNAs. Sample size: 120 (63 male, 57 female); Abbreviations: SkM, skeletal muscle; Amt, amount of exercise; Int, intensity of exercise. **Panel A** represents the total number of pathways for each database category (Gene Ontology-Biological process, Gene Ontology-Molecular Function, KEGG, and Reactome) derived from miEAA server (microRNA enrichment, annotation, and analysis) using all SkM miRNAs rank ordered based on estimate scores from highest to lowest. Significance was set at p<0.05. **Panels B** represents the top 20 amount and intensity pathways for each database category. Red dots represent enriched pathways whereas blue dots represent depleted; size of the dot represents the -log10 P-value. **Panel C** depicts the oxidative phosphorylation pathway with stars representing regions along the mitochondrial network where miRNAs are regulating target mRNAs; the 9 miRNAs identified in this pathway are predicted to regulate 31 mRNAs within the oxidative phosphorylation pathway. Pathway graphics were generated in R using Pathview.

In fully adjusted SkM models, exercise amount (accounting for intensity) modified 25 piRNAs (p<0.05) with 10 meeting Q<0.1 (**Table 2, Supplemental Table S4**). Computational analysis showed that these piRNAs had predicted interactions with several genes involved in metabolic processes (*NDUFS7, AMPD3, B3GALT1*, and *DHFR*). Of these piRNAs, DQ593358 was predicted to target the transfer RNA (tRNA) repeat element, tRNA-Asn-AAC (Asparaginyl-tRNA synthetase) involved in aminoacyl tRNA biosynthesis, which plays a crucial role in protein translation. Exercise intensity (accounting for amount) modified 72 piRNAs (p<0.05) with 3 meeting Q<0.1 (**Table 2, Supplemental Table S4**). Computational analysis showed that these piRNAs had predicted gene interactions related to ErbB signaling (*CBLC, ERBB4, PRKCB*, and *PTK2*), calcium signaling (*CACNA11, ERBB4, GRIN2A, PRKCB*, and *RYR2*), and MAPK Signaling (*MKNK1, MECOM, CACNA11, ERBB4, MAPT*, and *PRKCB*). Importantly, these results are computationally driven and are based on sequence complementarity, thus more work is needed to validate interactions.

**Table 2:**
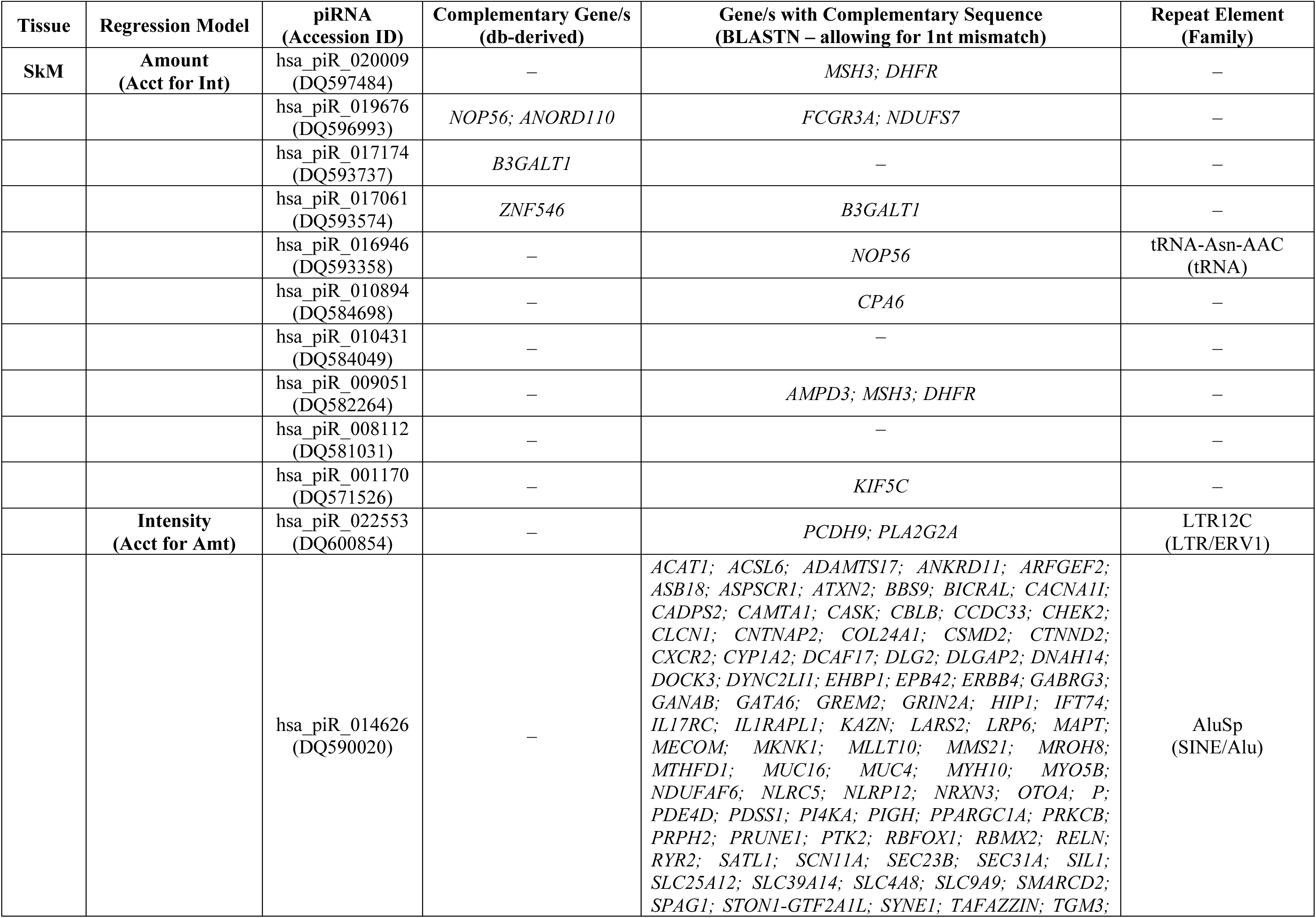
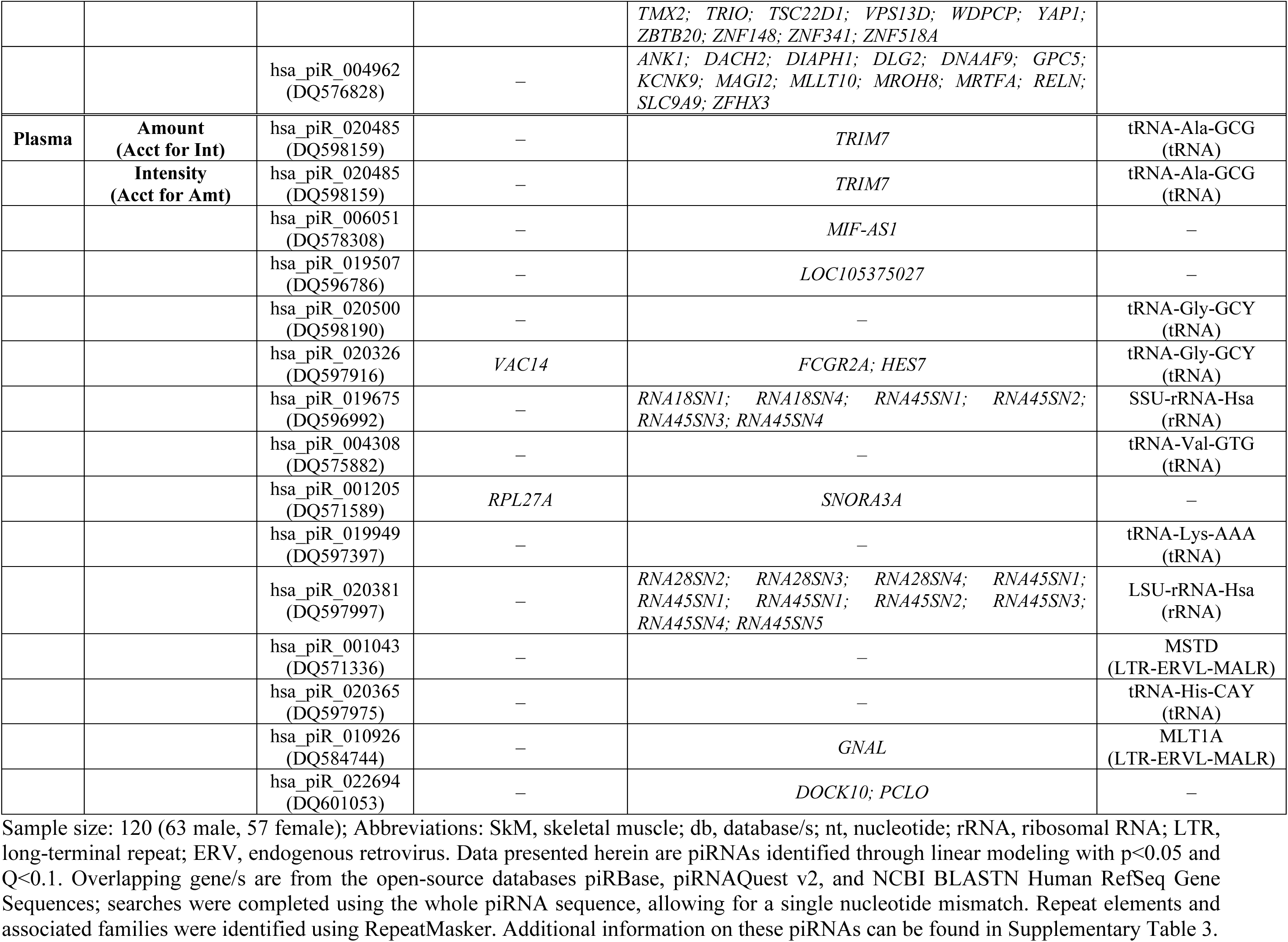
Significant SkM and Plasma piRNAs and Predicted mRNA Targets.

Taken together, our miRNA and piRNA findings, highlight dynamic effects of aerobic exercise on smRNA expression in SkM, converging on pathways involved in metabolic response, transcriptional and translational regulation, and cell cycle maintenance, thus emphasizing the role of epigenetic modifiers, such as smRNAs, in modulating exercise-induced adaptation.

### Effects of Aerobic Exercise on Plasma smRNA Expression

We also evaluated the effects of amount and intensity of exercise on circulating plasma smRNA expression. Similar to SkM modeling, we evaluated effects of amount—accounting for intensity, and intensity—accounting for amount, adjusting for baseline smRNA expression, age, race, sex, baseline BMI, and percent change in bodyweight, (**Figure 4a; Supplemental Table S5**).

**Figure 4:**
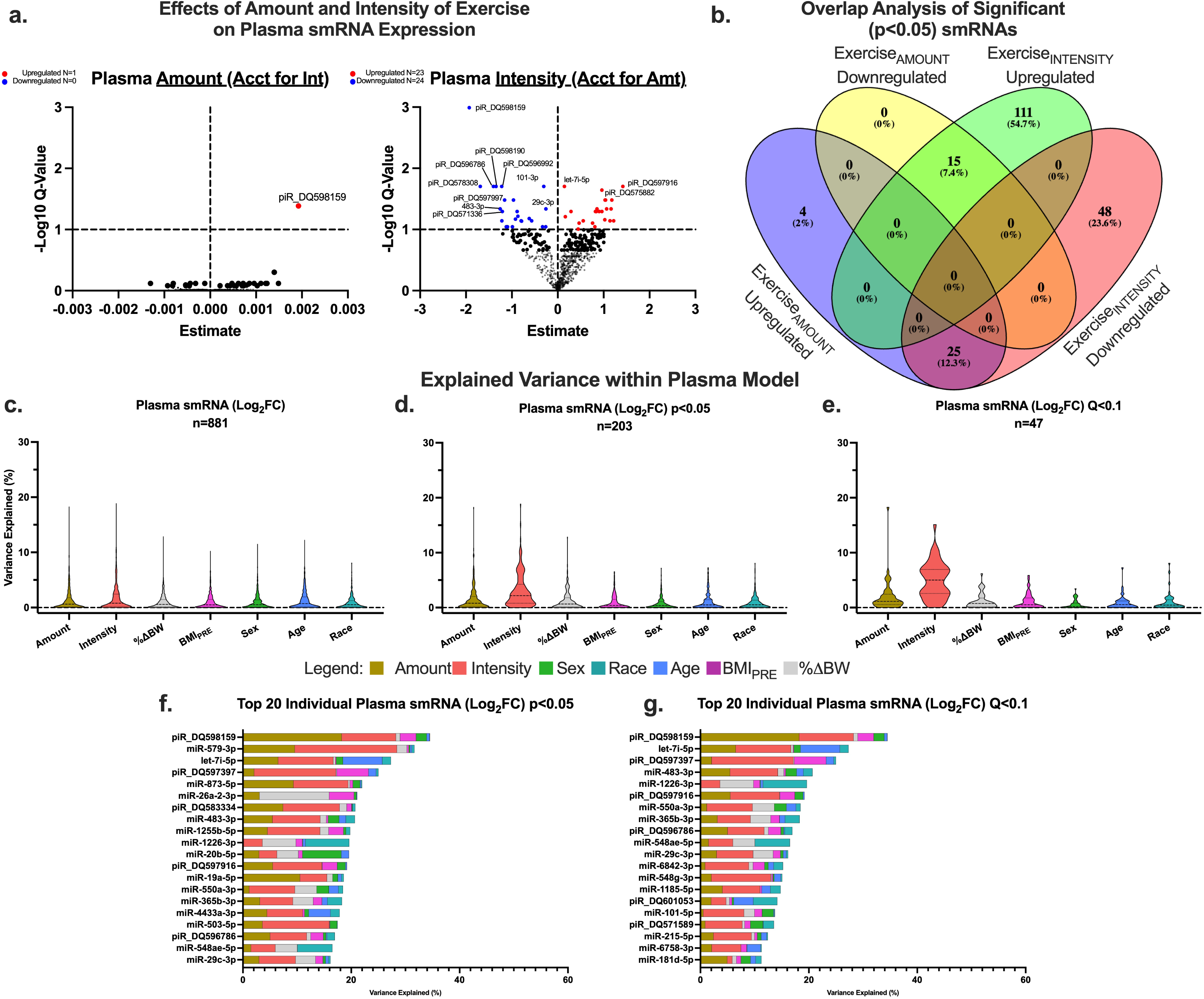
Effects of Exercise Amount and Intensity on smRNA Expression in Circulating Plasma. Sample size: 120 (63 male, 57 female); Abbreviations: plasma, circulating plasma; smRNA, small RNA; Amt, amount of exercise; Int, intensity of exercise; BMI_PRE_, Baseline BMI; %ΔBW, percent change in bodyweight. **Panel A** represents volcano plots of the -Log10 Q-value and estimate for adjusted plasma smRNA models. Non-significant smRNAs are depicted in black, significantly upregulated (p<0.05; Q<0.1) plasma smRNAs are depicted in red, and significantly (p<0.05; Q<0.1) downregulated plasma smRNAs are depicted in blue. **Panel B** represents analysis of overlap whereas the Venn diagram shows the total number of nominally significant (p<0.05) up-and down-regulated plasma smRNAs for amount of exercise (accounting for intensity) and for intensity of exercise (accounting for amount) and the specific plasma smRNAs that overlap in these variables. 25 smRNAs were identified as upregulated with exercise amount but downregulated with exercise intensity; 15 smRNAs were identified as downregulated by exercise amount but upregulated with exercise intensity. **Panel C-E** represent explainable variance with our linear modeling where violin plots depict analysis on all plasma smRNAs as well as plasma smRNAs with p<0.05 and Q<0.1. **Panel F-G** are bar plots depicting variance in the top 20 plasma smRNAs with p<0.05 and Q<0.1; the remaining variance not depicted is attributable to unmeasured factors accounted for in residuals.

We identified 44 ‘amount’-responsive smRNAs (p<0.05, 28 miRNA; 16 piRNA); only piRNA DQ598159 was significant and upregulated based on a Q<0.1 threshold. We identified 199 intensity-responsive smRNAs (p<0.05; 156 miRNA; 43 piRNA), with 47 (33 miRNA; 14 piRNA) meeting the Q<0.1 significance threshold (23 upregulated, 24 downregulated).

A total of 40 DE smRNAs (p<0.05) were independently associated with both exercise amount and intensity. Similar to SkM smRNA responses, in all cases, the effects of amount and intensity were divergent: 25 smRNAs were downregulated by intensity but upregulated by amount, whereas 15 showed the opposite pattern, being upregulated by intensity but downregulated by amount (**Figure 4b**). Of the smRNAs upregulated by exercise amount but downregulated by intensity (n=25), 14 were miRNAs and 11 were piRNAs. Conversely, those showing the opposite pattern, downregulated by amount and upregulated by intensity (n=15), were primarily miRNAs (n=10).

VariancePartition analysis showed that our model explained a mean 8.6% of the total variance in plasma smRNAs (n=881 smRNAs; **Figure 4c; Supplemental Table S6**). For exercise-responsive plasma smRNAs (p<0.05; **Figure 4d**), the explained variance increased to a mean 9.7% with exercise amount accounting for 1.5% (range: 0.0 to 18.2%) and exercise intensity 3.0% (range: 0.0 to 18.9%). Among the most significant exercise-responsive smRNAs (Q<0.1; **Figure 4e**), the model explained a mean 11.9% of variance, with exercise amount accounting for a mean 2.0 % (range: 0.0% to 18.2%) and intensity 5.0% (range: 0.0 to 15.1%). Notably, in top exercise-responsive plasma smRNAs (**Figure 4f-g**), intensity accounted for a greater share of variance than amount, in contrast to SkM, where amount predominated. This highlights a tissue-specific divergence in the regulation of smRNA expression by exercise parameters.

### Pathways Associated with Plasma smRNAs

Pathway analysis of plasma miRNAs identified 4,441 enriched or depleted pathways (p <0.05) across KEGG, Reactome, and GO libraries. Notably, exercise amount was predominantly associated with pathway enrichment (2,475 pathways), whereas exercise intensity was primarily associated with pathway depletion (1,872 pathways) (**Figure 5a–b; Supplemental Table S7**), highlighting opposing effects. Top pathways, ranked on p-value, were primarily related to metabolic regulation, cell cycle, and proteostasis. Notably, after network pruning, insulin signaling was enriched with exercise amount and depleted with intensity (p <0.001), with 13 miRNAs predicted to regulate 9 mRNAs within this pathway (**Figure 5c**).

**Figure 5:**
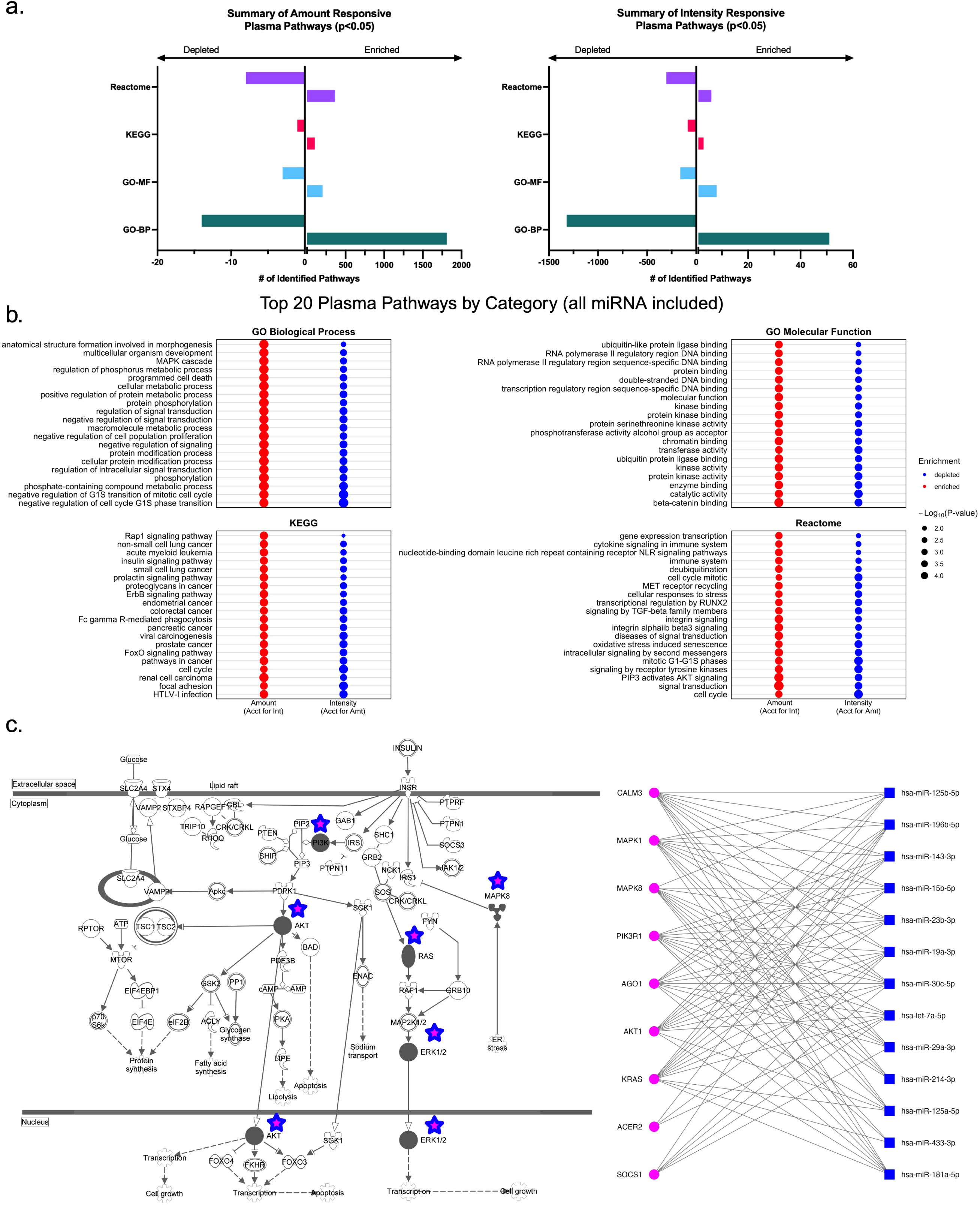
Pathways Associated with Plasma smRNAs. Sample size: 120 (63 male, 57 female); Abbreviations: plasma, circulating plasma; Amt, amount of exercise; Int, intensity of exercise. **Panel A** represents the total number of pathways for each database category (Gene Ontology-Biological process, Gene Ontology-Molecular Function, KEGG, and Reactome) derived from miEAA server (microRNA enrichment, annotation, and analysis) using all plasma miRNAs rank ordered based on estimate scores from highest to lowest. Significance was set at p<0.05. **Panels B** represents the top 20 amount and intensity pathways for each database category. Red dots represent enriched pathways whereas blue dots represent depleted; size of the dot represents the -log10 P-value. **Panel C** depicts the insulin signaling pathway with stars representing nodes within the pathway where miRNAs are regulating target mRNAs; 13 miRNAs were predicted to regulate 9 mRNAs within the insulin signaling pathway. Pathway graphics were generated in Ingenuity Pathway Analysis (IPA; Qiagen).

In fully adjusted plasma models, exercise amount was associated with changes in 16 piRNAs (p <0.05), though only one (DQ598159) remained significant after multiple testing correction Q<0.1 (**Table 2, Supplemental Table S3**). Computational analysis showed that this piRNA is predicted to target the gene *TRIM7* and a tRNA-Alanine repeat element. Exercise intensity altered 43 piRNAs, with 14 passing the Q <0.1 threshold (**Table 2, Supplemental Table S3**).

Computational analysis showed that these piRNAs are predicted to interact with genes involved in ribosome function (*RNA18SN1, RNA18SN4, RNA28SN2, RNA28SN3, RNA28SN4, RNA45SN3,* and *RNA45SN4*), and target various genomic repeats, including tRNAs, rRNAs, and long terminal repeats. Similar to our SkM findings, these results are computationally driven and require further investigation to validate interactions.

Taken together, these findings identify exercise-induced changes in plasma smRNA expression, with divergent enrichment signatures across metabolic and proteostatic pathways – including translational control, supporting their potential roles as both modulators of systemic health and biomarkers of exercise adaptation.

### Correlation of Plasma and Skeletal Muscle smRNA

Given the exercise responses of many SkM and plasma smRNAs, we performed pairwise Spearman correlation analyses to explore potential relationships between smRNA expression in the two tissues. We identified significant correlations (p<0.05; Spearman *ρ* range: -0.261 to 0.293) of fold changes in SkM and plasma for 36 smRNAs, 23 positively and 13 negatively correlated (**Supplemental Table S8**). The strongest positive correlations were among miRs -371a-5p (ρ=0.293) and -106b-5p (ρ=0.261) involved in proteostasis and cell cycle maintenance, and piR DQ595023 (ρ=0.266) involved in proteostasis through targeting of *EIF5*; the strongest negative correlations were observed for miRs -34c-5p (ρ=-0.261), -210-3p (ρ=-0.238), and -450a-2-3p (ρ=0.230) involved with cell cycle maintenance. Despite small but modest correlations, this analysis suggests potential for coordinated smRNA responses between SkM and plasma following exercise.

### Association of Exercise Responsive smRNA and Change in Insulin Sensitivity Index

In a subset of our cohort with insulin sensitivity index (IS_i_) data (n=96), IS_i_ increased a mean 28.7%. Not unexpectedly, most participants (n=61) showed improvement (mean +64.2%), while the remaining participants (n=35) showed either no change or a decrease in IS_i_ (mean -33.2%) (**Supplemental Figure S2**). Using univariable linear regression, we identified percent change in bodyweight and smRNAs from both SkM and plasma associated with change in IS_i_ (**Table 3; Supplemental Table S9)**. In SkM models, the change of 9 smRNAs were associated with change in IS_i_ (p<0.05); of these, 6 smRNAs (miRs -1246, 29b-1-5p, 131-3p, and 376a-2-5p; piRs DQ582536 and DQ596805) exhibited exercise-induced expression changes aligned with IS_i_ improvement. In plasma models, the change of 12 smRNAs were associated with change in IS_i_ (p<0.05), with 5 (miRs -365b-3p, -362-3p, -210-3p, -365a-3p, and -3688-3p) showing exercise-induced changes consistent with improved IS_i_. As expected, greater reductions in bodyweight were associated with improvements (increases) in IS_i_. While exploratory, these findings highlight distinct sets of exercise-responsive SkM- and plasma-derived smRNAs associated with exercise-induced improvements in insulin sensitivity, yielding potential SkM-specific regulators and circulating biomarkers of metabolic response to exercise.

**Table 3:** Statistically Significant Univariable Associations of Change in smRNA with Change in in Sensitivity Index.

|  | Exercise Effects |  |  |  |  |  |
| --- | --- | --- | --- | --- | --- | --- |
| Variable | Amount | Intensity | Std. Beta (95% CI) | R<sup>2</sup> | P= | Q= |
| **<u>SkM Variables</u>** |  |  |  |  |  |  |
| miR-7-1-3p |  | ↓ | 0.339 (0.149, 0.530) | 0.115 | 0.001 | 0.248 |
| piR_DQ582536 |  | ↑ | 0.261 (0.066, 0.456) | 0.068 | 0.010 | 0.989 |
| miR-378h |  | ↓ | 0.216 (0.018, 0.413) | 0.047 | 0.035 | 0.989 |
| miR-1246 | ↓ | ↑ | 0.215 (0.018, 0.412) | 0.046 | 0.035 | 0.989 |
| miR-29b-1-5p | ↑ | ↓ | -0.215 (-0.412, -0.017) | 0.046 | 0.036 | 0.989 |
| miR-191-3p | ↑ |  | 0.211 (0.013, 0.408) | 0.044 | 0.039 | 0.989 |
| miR-376a-2-5p | ↑ | ↓ | 0.209 (0.012, 0.407) | 0.044 | 0.041 | 0.989 |
| piR_DQ596805 |  | ↑ | 0.206 (0.008, 0.404) | 0.042 | 0.044 | 0.989 |
| miR-1297 |  | ↓ | 0.201 (0.003, 0.399) | 0.040 | 0.050 | 0.989 |
| **<u>Plasma Variables</u>** |  |  |  |  |  |  |
| miR-505-3p | ↑ |  | -0.352 (-0.541, -0.162) | 0.124 | <0.001 | 0.092 |
| miR-365b-3p | ↑ | ↓ | -0.299 (-0.492, -0.106) | 0.089 | 0.003 | 0.324 |
| miR-362-3p | ↑ | ↓ | -0.265 (-0.460, -0.070) | 0.070 | 0.009 | 0.447 |
| miR-598-3p |  | ↑ | -0.257 (-0.453, -0.062) | 0.066 | 0.011 | 0.447 |
| let-7b-3p | ↑ | ↓ | -0.253 (-0.448, -0.057) | 0.064 | 0.013 | 0.447 |
| miR-210-3p |  | ↓ | -0.249 (-0.445, -0.054) | 0.062 | 0.014 | 0.447 |
| miR-942-5p |  | ↑ | -0.248 (-0.443, -0.052) | 0.061 | 0.015 | 0.447 |
| miR-133a-3p |  | ↑ | -0.240 (-0.436, -0.043) | 0.057 | 0.019 | 0.486 |
| miR-145-3p | ↑ |  | -0.232 (-0.429, -0.036) | 0.054 | 0.023 | 0.528 |
| miR-365a-3p |  | ↓ | -0.216 (-0.413, -0.018) | 0.047 | 0.035 | 0.678 |
| miR-6818-3p | ↓ | ↑ | -0.214 (-0.412, -0.017) | 0.046 | 0.036 | 0.678 |
| miR-3688-3p |  | ↓ | -0.208 (-0.406, -0.011) | 0.043 | 0.042 | 0.678 |

Sample size: 96 (52 male, 44 female); Abbreviations: SkM, skeletal muscle; smRNA, small RNA; CI, confidence interval; miR, microRNA; piR\_DQ, piwi-interacting RNA accession ID. Data presented herein represent variables associated with change in insulin sensitivity index (IS<sub>i</sub>), identified via univariable linear regression modeling. Data are presented as standardized estimates with 95% CIs, R<sup>2</sup>, p-value, and Benjamini-Hochberg corrected Q-value. Significant (p<0.05) up- or down-regulation of smRNAs by exercise amount and intensity are indicated by upward and downward arrows; black arrows indicate an exercise-induced change in smRNA consistent with improved IS<sub>i</sub>. Change in plasma miRs -365b-3p and 942-5p association with change in IS<sub>i</sub> were significant at both p<0.05 and Q<0.1.

## DISCUSSION

Exercise amount and intensity exhibited an inverse relationship on smRNA that reflects the inverse relationship observed at a physiological level. As intensity increases relative to ̇VO_2_max, metabolic efficiency modestly declines due to greater reliance on less efficient type II muscle fibers and anaerobic energy pathways^51,52^; this is accompanied by a substrate shift from fat oxidation to carbohydrate metabolism^53^. These physiological adaptations are likely mediated, in part, by smRNA signaling within SkM and from SkM to other organ systems. Our study used untargeted smRNA sequencing to examine how an 8–9-month aerobic exercise intervention, varying in amount and intensity, altered smRNA expression profiles. This comprehensive approach captured data on thousands of miRNAs and piRNAs – a class of smRNAs whose exercise-induced modifications in somatic cells had not been previously reported. Our data show that exercise amount predominantly influences SkM smRNA expression, whereas exercise intensity primarily affects plasma smRNA profiles. Pathway analyses via miEAA using the GSEA algorithm and all miRNAs, identified coordinated targeting patterns across key metabolic, transcriptional, translational, and cell cycle pathways, including oxidative phosphorylation and insulin signaling. Importantly, we identified 6 SkM and 5 plasma smRNAs, whose exercise-induced change, were consistent with improvements in IS_i_ following chronic aerobic exercise. While previous studies have extensively investigated miRNA changes in SkM and plasma in response to exercise^3,25,54^, this study is the first to concurrently examine the distinct roles of exercise amount and intensity on smRNA profiles.

A major strength of this study is the use of untargeted smRNA sequencing, enabling detection of 3,163 smRNAs in SkM (2,558 miRNAs, 605 piRNAs) and 2,801 smRNAs in plasma (2,544 miRNAs, 257 piRNAs). While numerous studies have explored miRNA regulation through exercise^55–58^, they typically focused on fewer than ten miRNAs using RT-qPCR or limited arrays^59^, leaving gaps in the broader understanding of smRNA expression. Our analysis identified 139 significant smRNAs (p<0.05, Q<0.1) in SkM, most (∼90%) of which were responsive to exercise amount. In plasma, 48 significant smRNAs were identified (p<0.05, Q<0.1), with ∼98% responsive to exercise intensity. Of these, only ∼15 smRNAs have been reported in prior exercise literature^3,25^.

Consistent with prior exercise literature^56,57,60^, within SkM, exercise amount significantly upregulated classical myomiRs including miRs -1-5p and -133a-3p, and downregulated miRs -1-3p and -206. We also observed that aerobic exercise upregulated miRs -153-3p, -296-5p, -483-3p, -488-3p, and the miR-30 family, all implicated in muscle regeneration, growth, and metabolism. Differential expression of the miR-30 family has been linked to physiological cardiac hypertrophy – a hallmark of chronic aerobic exercise adaptation^61–66^. Increased expression of miRs -153-3p, - 296-5p, -483-3p, and -488-3p is associated with mitochondrial biogenesis, lipid regulation, inhibition of myoblast proliferation, and enhanced metabolic capacity^67–73^. Increased miR-483-3p expression is also associated with muscle atrophy in humans and mice^74^.

In plasma, the 48 significant smRNAs (33 miRNAs and 14 piRNAs) were responsive to exercise intensity, with one piRNA (DQ598159) exhibiting dual responsiveness, upregulated by exercise intensity and downregulated by exercise amount. Several miRNAs, including miRs -132-5p, -181d-5p, -424-3p, -589-3p, and -1226-3p have prior associations to exercise, inflammation^75–82^, or metabolic regulation^83^, and five additional miRNAs from our data (e.g., -210-3p, -365a/b-3p, -3688-3p, -362-3p) showed changes consistent with improved insulin signaling. Notably, some exhibited dual responses, being upregulated by exercise amount but downregulated by intensity. Circulating plasma miRNAs are stabilized within extracellular vesicles, HDL particles, and Argonaute2-bound complexes^26,84–86^, and experimental systems demonstrate their capacity to modulate gene expression in recipient cells. While these mechanisms support the plausibility of intercellular miRNA communication, we interpret plasma smRNAs here as circulating molecular signatures associated with exercise dose, stress-response, and metabolic adaptation rather than direct evidence of endocrine target regulation in skeletal muscle.

After low-expression filtering, the median raw read count per smRNA per library in SkM was 19.0 at PRE and 20.0 at POST. In plasma, the corresponding per-library median raw count was 0 at both time points. With the exception of one SkM smRNA, the counts of SkM smRNAs associated with changes in insulin sensitivity were similar to the overall median counts per participant (19 counts pre-exercise and 20 counts post-exercise) across the N=1188 distinct smRNAs detected. In contrast, except for one smRNA, the plasma smRNAs associated with changes in insulin sensitivity index had counts above the overall median (range 23-273) for all detected smRNAs (n=881). Canonical miRNA function involves Argonaute (AGO)/RNA-induced silencing complex (RISC)-mediated repression of target transcripts; however, biological activity is not limited to this mechanism nor solely determined by miRNA abundance. AGO-mediated repression is governed by stoichiometric relationships among miRNA abundance, target-site availability, and effective RISC capacity rather than simple rank abundance thresholds^87,88^. Moreover, miRNA regulation operates within a many-to-many framework in which individual miRNAs influence numerous transcripts and individual transcripts are also subject to combinatorial control^89–91^, typically producing modest, distributed network-level effects rather than dominance by the most abundant miRNAs^92,93^. Accordingly, our pathway analyses are intended to provide network-level biological context rather than infer direct repression of specific targets *in vivo*.

Pathway analysis identified approximately 1,300 SkM and 4,400 plasma miRNA-associated pathways, with enrichment in metabolic regulation, transcriptional and translational control, and cell cycle processes. In SkM, oxidative phosphorylation emerged as a top pathway, underscoring its essential role in sustaining ATP production during prolonged aerobic exercise^94–96^. Additionally, enrichment of glucose-6-phosphate metabolism and carbohydrate kinase activity highlights the importance of glycolytic flux, particularly hexokinase-mediated glucose phosphorylation^53^. Insulin signaling–related pathways were enriched in both SkM and plasma, supporting a coordinated role for smRNAs in exercise-mediated glucose metabolism.

We identified six SkM and five plasma exercise-responsive smRNAs whose modulation was associated with improvements in IS_i_. In SkM these included: miR-191-3p and miR-376a-2-5p (upregulated by amount); miR-1246, piRNAs DQ5825356 and DQ596805 (upregulated by intensity); and miR-29b-1-5p (downregulated by intensity). Downregulation of miR-29b-1-5p is consistent with evidence showing that inhibition of the miR-29 family improves insulin-stimulated glucose uptake and insulin sensitivity^97,98^ and enhances *GLUT4* transcription in skeletal muscle^99,100^. These findings provide mechanistic context for miR-191-3p, which has been positively associated with fasting insulin and HOMA-IR in humans^101^ and shown to impair GLUT4 translocation and glucose uptake^102^.

In plasma, exercise-responsive smRNA associated with improvements in ISi included: miRs-210-3p, -365a-3p, -3688-3p, -365b-3p, and -362-3p (downregulated by intensity). Prior studies implicate miR-365a/b-3p in diabetes risk^103,104^ and show that miR-210-3p suppresses insulin sensitivity via NFκB signaling and impaired GLUT4 translocation^105,106^. Additionally, in our cohort, modulation of SkM miRs by exercise (−30d-5p downregulated by amount but upregulated by intensity; -26b-5p, -27a-3p, -29a-5p downregulated by amount), yielded changes consistent with improved IS_i_, based on literature showing that insulin sensitivity has a complicated dose-response relationship with exercise^35,107^, mediated through PI3K/AKT-dependent GLUT4 translocation downstream of insulin receptor activation^10,108,109^. Multiple miRNAs, including miR-15b, -26b, -27a, -29a, -30d, -106b, -195, and -424-5p, regulate this pathway^109–112^. Collectively, these findings indicate that exercise-responsive smRNAs in SkM and plasma converge on shared regulatory nodes controlling insulin signaling and glucose transport, particularly via GLUT4 and PI3K/AKT signaling.

Data on differential expression profiles of piRNAs with various interventions are sparse. Functionally, piRNAs silence transposable elements (TEs) and safeguard germline DNA^113–116^. Our data complement emerging evidence that piRNAs also regulate somatic cell gene expression based on sequence complementarity to mRNAs^117–119^, supporting broader biological relevance for this class of smRNAs outside of germline cells^22^. To date, only one study has evaluated the effects of exercise on piRNA expression, reporting that chronic aerobic exercise downregulated piRNAs DQ597945 and DQ571511 in human spermatozoa^120^. We observed a nominal downregulation of DQ597945 in plasma while DQ571511 was nominally upregulated in SkM. Importantly, our piRNA-to-mRNA and piRNA-to-genomic repeat interactions rely on computational algorithms and have yet to be experimentally validated, yielding exciting new directions in smRNA biology worth investigating.

Our study has several strengths, including being among the first to apply untargeted smRNA deep sequencing in a human cohort undergoing a controlled, chronic, and closely monitored exercise intervention. The trial design enabled us to examine the independent effects of exercise amount and intensity on smRNA (miRNA and piRNA) expression profiles in both SkM and plasma. Our investigation leveraged 120 participants with matched SkM and plasma samples, providing adequate power for regression modeling. Although the relatively small number of non-exercise controls (n=11) limits precision at the lowest end of the exercise intensity spectrum, exercise amount and intensity were modeled as continuous variables, for which statistical power depends primarily on overall sample size and predictor variance^121,122^. Accordingly, the study was adequately powered to detect smRNAs associated with exercise amount and intensity. With respect to smRNAs, current prediction of smRNA-to-mRNA, particularly piRNA-to-mRNA interactions, have some limitations: functional interactions are not always a result of perfect sequence alignment, and only a subset of identified smRNAs have been experimentally validated, leaving accurate prediction of mRNA interactions as an unresolved experimental and computational challenge ^123^. Although numerous miRNAs have been identified over the past three decades, many remain experimentally unvalidated^124,125^, highlighting the necessity of confirming the novel smRNAs identified in this study. This issue underscores the importance of further validating novel smRNA sequences and functions to clarify their roles in exercise-induced regulatory pathways.

In summary, exercise amount was the primary modulator of skeletal muscle smRNA expression with the majority of exercise-responsive smRNAs being miRNA. Conversely, exercise intensity predominantly modulated plasma smRNA expression. Our analysis of explainable variance showed exercise amount accounted for the largest proportion of variance in SkM smRNA whereas intensity accounted for the largest proportion in plasma smRNA, confirming the trend we observed in our linear regression analysis. Top pathways influenced by SkM and plasma miRNAs were related to metabolism, transcriptional and translational regulation, and cell cycle maintenance. We identified 11 smRNAs (6 in SkM and 5 in plasma) whose exercise-induced changes were consistent with improvements in IS_i_. These adaptations may contribute to inter-individual variability in metabolic responses, with both beneficial and detrimental effects, and highlight novel targets influencing insulin secretion and glucose uptake in skeletal muscle. Future work will test their efficacy within the insulin signaling cascade to assess therapeutic potential in individuals unable or unwilling to exercise. Collectively, our findings provide strong evidence that exercise alters smRNA expression in both plasma and skeletal muscle and reveal candidate regulators of metabolic and other health-related outcomes.

## Additional Information

### Data availability statement

Reasonable requests for data and resources should be directed to Dr Christopher Vann, Dr William Kraus, and Dr Virginia Kraus. Upon publication, processed sequencing data will be available for download via synapse.org (accession ID: syn68234751) at https://doi.org/10.7303/syn68234751. These data are comprised of the raw and log_2_CPM counts for SkM and plasma smRNA. Supplemental Tables S1-S11 and Supplemental Figures S1 and S2 are available via FigShare at: https://doi.org/10.6084/m9.figshare.31018321.

## Ethics statement

The STRRIDE trials that generated the tissue and clinical data used herein were reviewed by the Duke University and East Carolina University Institutional Review Boards. Participants provided oral and written informed consent to participate in this study.

## Author contributions

CGV, VBK, WEK, CFA, SM, KMH, and MJH conceived the study. CGV and MCO performed wet laboratory experiments. WCB, LMR, MJH, WEK, KMH, and CGV curated the data. CGV, AB, SM, and SIN performed the statistical analyses. VBK, WEK, and CFA acquired funding for this study. CGV, VBK, WEK, CFA, SM, AB, SIN, and MCO devised the methodologies for the study. CGV, VBK, and WEK were responsible for overall project supervision and drafting of the manuscript. All authors read, edited, and approved the final version of the manuscript.

## Acknowledgements

The authors would like to thank Dr. Nicolas Devos, Dr. Devi Swain Lenz, Laura-Leigh Rowlette, and the Duke University Sequencing and Genomics Technology Core for performing smRNA sequencing.

## Abstract Figure

Abbreviations: Amt, exercise amount; Int, exercise intensity; PRE, biological sample collection prior to intervention; POST, biological sample collection following intervention; smRNA, small RNA; BMI, body mass index; SkM, skeletal muscle. Graphical Abstract was generated using BioRender.com.

## REFERENCES

1 Guo, B. et al. Molecular Communication from Skeletal Muscle to Bone: A Review for Muscle-Derived Myokines Regulating Bone Metabolism. Calcif Tissue Int 100, 184–192 (2017). 10.1007/s00223-016-0209-4

2 Plomgaard, P., Halban, P. A. & Bouzakri, K. Bimodal impact of skeletal muscle on pancreatic β-cell function in health and disease. Diabetes Obes Metab 14 **Suppl 3**, 78–84 (2012). 10.1111/j.1463-1326.2012.01641.x

3 Silva, G. J. J., Bye, A., El Azzouzi, H. & Wisloff, U. MicroRNAs as Important Regulators of Exercise Adaptation. Prog Cardiovasc Dis 60, 130–151 (2017). 10.1016/j.pcad.2017.06.003

4 Aronoff, S. L., Berkowitz, K., Shreiner, B. & Want, L. Glucose Metabolism and Regulation: Beyond Insulin and Glucagon. Diabetes Spectrum 17, 183–190 (2004). 10.2337/diaspect.17.3.183

5 Brown, M. D., Moore, G. E., Korytkowski, M. T., McCole, S. D. & Hagberg, J. M. Improvement of insulin sensitivity by short-term exercise training in hypertensive African American women. Hypertension 30, 1549–1553 (1997). 10.1161/01.hyp.30.6.1549

6 Kahn, S. E. et al. Effect of exercise on insulin action, glucose tolerance, and insulin secretion in aging. Am J Physiol 258, E937–943 (1990). 10.1152/ajpendo.1990.258.6.E937

7 Kraus, W. E. et al. Studies of a targeted risk reduction intervention through defined exercise (STRRIDE). Med Sci Sports Exerc 33, 1774–1784 (2001). 10.1097/00005768-200110000-00025

8 Bergman, R. N., Finegood, D. T. & Ader, M. Assessment of Insulin Sensitivity in Vivo*. Endocrine Reviews 6, 45–86 (1985). 10.1210/edrv-6-1-45

9 Slentz, C. A. et al. Effects of exercise training alone vs a combined exercise and nutritional lifestyle intervention on glucose homeostasis in prediabetic individuals: a randomised controlled trial. Diabetologia 59, 2088–2098 (2016). 10.1007/s00125-016-4051-z

10 Nigi, L. et al. MicroRNAs as Regulators of Insulin Signaling: Research Updates and Potential Therapeutic Perspectives in Type 2 Diabetes. Int J Mol Sci 19 (2018). 10.3390/ijms19123705

11 Jacques, M. et al. Epigenetic changes in healthy human skeletal muscle following exercise-a systematic review. Epigenetics 14, 633–648 (2019). 10.1080/15592294.2019.1614416

12 Lacal, I. & Ventura, R. Epigenetic Inheritance: Concepts, Mechanisms and Perspectives. Front Mol Neurosci 11, 292 (2018). 10.3389/fnmol.2018.00292

13 O’Brien, J., Hayder, H., Zayed, Y. & Peng, C. Overview of MicroRNA Biogenesis, Mechanisms of Actions, and Circulation. Front Endocrinol (Lausanne) 9, 402 (2018). 10.3389/fendo.2018.00402

14 Chekulaeva, M. & Filipowicz, W. Mechanisms of miRNA-mediated post-transcriptional regulation in animal cells. Curr Opin Cell Biol 21, 452–460 (2009). 10.1016/j.ceb.2009.04.009

15 Friedman, R. C., Farh, K. K., Burge, C. B. & Bartel, D. P. Most mammalian mRNAs are conserved targets of microRNAs. Genome Res 19, 92–105 (2009). 10.1101/gr.082701.108

16 Wang, C. & Lin, H. Roles of piRNAs in transposon and pseudogene regulation of germline mRNAs and lncRNAs. Genome Biol 22, 27 (2021). 10.1186/s13059-020-02221-x

17 Zhang, F. & Wang, D. The Pattern of microRNA Binding Site Distribution. Genes (Basel) 8 (2017). 10.3390/genes8110296

18 Geles, K. et al. WIND (Workflow for pIRNAs aNd beyonD): a strategy for in-depth analysis of small RNA-seq data. F1000Res 10, 1 (2021). 10.12688/f1000research.27868.3

19 Yu, T. et al. The piRNA Response to Retroviral Invasion of the Koala Genome. Cell 179, 632–643 e612 (2019). 10.1016/j.cell.2019.09.002

20 Aravin, A. et al. A novel class of small RNAs bind to MILI protein in mouse testes. Nature 442, 203–207 (2006). 10.1038/nature04916

21 Haase, A. D. et al. PIWI-interacting RNAs: who, what, when, where, why, and how. Embo j 43, 5335–5339 (2024). 10.1038/s44318-024-00253-8

22 Kraus, V. B. et al. Select Small Non-Coding RNAs Are Determinants of Survival in Older Adults. Aging Cell 25, e70403 (2026). 10.1111/acel.70403

23 McCarthy, J. J. The MyomiR network in skeletal muscle plasticity. Exerc Sport Sci Rev 39, 150–154 (2011). 10.1097/JES.0b013e31821c01e1

24 Nielsen, S. et al. Muscle specific microRNAs are regulated by endurance exercise in human skeletal muscle. J Physiol 588, 4029–4037 (2010). 10.1113/jphysiol.2010.189860

25 Silva, F. C. D. et al. Effects of Physical Exercise on the Expression of MicroRNAs: A Systematic Review. J Strength Cond Res 34, 270–280 (2020). 10.1519/jsc.0000000000003103

26 Vann, C. G. et al. Differential microRNA profiles of intramuscular and secreted extracellular vesicles in human tissue-engineered muscle. Front Physiol 13, 937899 (2022). 10.3389/fphys.2022.937899

27 Wasfy, M. M. & Lee, I. M. Examining the Dose-Response Relationship between Physical Activity and Health Outcomes. NEJM Evid 1, EVIDra2200190 (2022). 10.1056/EVIDra2200190

28 Bateman, L. A. et al. Comparison of aerobic versus resistance exercise training effects on metabolic syndrome (from the Studies of a Targeted Risk Reduction Intervention Through Defined Exercise - STRRIDE-AT/RT). Am J Cardiol 108, 838–844 (2011). 10.1016/j.amjcard.2011.04.037

29 Hittel, D. S., Kraus, W. E., Tanner, C. J., Houmard, J. A. & Hoffman, E. P. Exercise training increases electron and substrate shuttling proteins in muscle of overweight men and women with the metabolic syndrome. J Appl Physiol (1985) 98, 168–179 (2005). 10.1152/japplphysiol.00331.2004

30 Ross, L. M. et al. Effects of Amount, Intensity, and Mode of Exercise Training on Insulin Resistance and Type 2 Diabetes Risk in the STRRIDE Randomized Trials. Front Physiol 12, 626142 (2021). 10.3389/fphys.2021.626142

31 Slentz, C. A. et al. Inactivity, exercise, and visceral fat. STRRIDE: a randomized, controlled study of exercise intensity and amount. J Appl Physiol (1985) 99, 1613–1618 (2005). 10.1152/japplphysiol.00124.2005

32 Slentz, C. A. et al. Effects of the amount of exercise on body weight, body composition, and measures of central obesity: STRRIDE--a randomized controlled study. Arch Intern Med 164, 31–39 (2004). 10.1001/archinte.164.1.31

33 Slentz, C. A. et al. Inactivity, exercise training and detraining, and plasma lipoproteins. STRRIDE: a randomized, controlled study of exercise intensity and amount. J Appl Physiol (1985) 103, 432–442 (2007). 10.1152/japplphysiol.01314.2006

34 Kraus, W. E. et al. Effects of the amount and intensity of exercise on plasma lipoproteins. N Engl J Med 347, 1483–1492 (2002). 10.1056/NEJMoa020194

35 Houmard, J. A. et al. Effect of the volume and intensity of exercise training on insulin sensitivity. J Appl Physiol (1985) 96, 101–106 (2004). 10.1152/japplphysiol.00707.2003

36 Slentz, C. A. et al. Effects of aerobic vs. resistance training on visceral and liver fat stores, liver enzymes, and insulin resistance by HOMA in overweight adults from STRRIDE AT/RT. Am J Physiol Endocrinol Metab 301, E1033–1039 (2011). 10.1152/ajpendo.00291.2011

37 Barberio, M. D. et al. Pyruvate Dehydrogenase Phosphatase Regulatory Gene Expression Correlates with Exercise Training Insulin Sensitivity Changes. Med Sci Sports Exerc 48, 2387–2397 (2016). 10.1249/mss.0000000000001041

38 Hoffman, G. E. & Schadt, E. E. variancePartition: interpreting drivers of variation in complex gene expression studies. BMC Bioinformatics 17, 483 (2016). 10.1186/s12859-016-1323-z

39 Coffman, C. J., Edelman, D. & Woolson, R. F. To condition or not condition? Analysing ’change’ in longitudinal randomised controlled trials. BMJ Open 6, e013096 (2016). 10.1136/bmjopen-2016-013096

40 Vickers, A. J. & Altman, D. G. Statistics notes: Analysing controlled trials with baseline and follow up measurements. BMJ 323, 1123–1124 (2001). 10.1136/bmj.323.7321.1123

41 Benjamini, Y. & Hochberg, Y. Controlling the False Discovery Rate: A Practical and Powerful Approach to Multiple Testing. Journal of the Royal Statistical Society. Series B (Methodological) 57, 289–300 (1995).

42 Aparicio-Puerta, E. et al. miEAA 2023: updates, new functional microRNA sets and improved enrichment visualizations. Nucleic Acids Res 51, W319–W325 (2023). 10.1093/nar/gkad392

43 Kern, F. et al. miEAA 2.0: integrating multi-species microRNA enrichment analysis and workflow management systems. Nucleic Acids Res 48, W521–w528 (2020). 10.1093/nar/gkaa309

44 Chang, L. & Xia, J. MicroRNA Regulatory Network Analysis Using miRNet 2.0. Methods Mol Biol 2594, 185–204 (2023). 10.1007/978-1-0716-2815-7_14

45 Chirn, G. W. et al. Conserved piRNA Expression from a Distinct Set of piRNA Cluster Loci in Eutherian Mammals. PLoS Genet 11, e1005652 (2015). 10.1371/journal.pgen.1005652

46 Ghosh, B. et al. piRNAQuest V.2: an updated resource for searching through the piRNAome of multiple species. RNA Biol 19, 12–25 (2022). 10.1080/15476286.2021.2010960

47. RepeatMasker Open-4.0 (2013).

48 Wang, J. et al. piRBase: a comprehensive database of piRNA sequences. Nucleic Acids Res 47, D175–D180 (2019). 10.1093/nar/gky1043

49 Sherman, B. T. et al. DAVID: a web server for functional enrichment analysis and functional annotation of gene lists (2021 update). Nucleic Acids Res 50, W216–W221 (2022). 10.1093/nar/gkac194

50 Wickham, H. ggplot2: Elegant Graphics for Data Analysis. 2 edn, (Springer Nature, 2016).

51 Hunter, G. R., Newcomer, B. R., Larson-Meyer, D. E., Bamman, M. M. & Weinsier, R. L. Muscle metabolic economy is inversely related to exercise intensity and type II myofiber distribution. Muscle Nerve 24, 654–661 (2001). 10.1002/mus.1051

52 Hunter, G. R., Weinsier, R. L., Bamman, M. M. & Larson, D. E. A role for high intensity exercise on energy balance and weight control. International Journal of Obesity 22, 489–493 (1998). 10.1038/sj.ijo.0800629

53 Hargreaves, M. & Spriet, L. L. Skeletal muscle energy metabolism during exercise. Nature Metabolism 2, 817–828 (2020). 10.1038/s42255-020-0251-4

54 Ultimo, S. et al. Influence of physical exercise on microRNAs in skeletal muscle regeneration, aging and diseases. Oncotarget 9, 17220–17237 (2018). 10.18632/oncotarget.24991

55 Margolis, L. M. et al. Upregulation of circulating myomiR following short-term energy restriction is inversely associated with whole body protein synthesis. Am J Physiol Regul Integr Comp Physiol 313, R298–R304 (2017). 10.1152/ajpregu.00054.2017

56 Nielsen, S. et al. Muscle specific microRNAs are regulated by endurance exercise in human skeletal muscle. J Physiol 588, 4029–4037 (2010). 10.1113/jphysiol.2010.189860

57 Russell, A. P. et al. Regulation of miRNAs in human skeletal muscle following acute endurance exercise and short-term endurance training. J Physiol 591, 4637–4653 (2013). 10.1113/jphysiol.2013.255695

58 van Rooij, E. et al. A family of microRNAs encoded by myosin genes governs myosin expression and muscle performance. Dev Cell 17, 662–673 (2009). 10.1016/j.devcel.2009.10.013

59 Sapp, R. M., Shill, D. D., Roth, S. M. & Hagberg, J. M. Circulating microRNAs in acute and chronic exercise: more than mere biomarkers. J Appl Physiol (1985) 122, 702–717 (2017). 10.1152/japplphysiol.00982.2016

60 Drummond, M. J., McCarthy, J. J., Fry, C. S., Esser, K. A. & Rasmussen, B. B. Aging differentially affects human skeletal muscle microRNA expression at rest and after an anabolic stimulus of resistance exercise and essential amino acids. Am J Physiol Endocrinol Metab 295, E1333–1340 (2008). 10.1152/ajpendo.90562.2008

61 Backes, C. et al. Blood born miRNAs signatures that can serve as disease specific biomarkers are not significantly affected by overall fitness and exercise. PLoS One 9, e102183 (2014). 10.1371/journal.pone.0102183

62 Li, J. et al. Mir-30d Regulates Cardiac Remodeling by Intracellular and Paracrine Signaling. Circ Res 128, e1–e23 (2021). 10.1161/CIRCRESAHA.120.317244

63 Melman, Y. F. et al. Circulating MicroRNA-30d Is Associated With Response to Cardiac Resynchronization Therapy in Heart Failure and Regulates Cardiomyocyte Apoptosis: A Translational Pilot Study. Circulation 131, 2202–2216 (2015). 10.1161/CIRCULATIONAHA.114.013220

64 Ogasawara, R. et al. MicroRNA expression profiling in skeletal muscle reveals different regulatory patterns in high and low responders to resistance training. Physiol Genomics 48, 320–324 (2016). 10.1152/physiolgenomics.00124.2015

65 Radom-Aizik, S., Zaldivar, F., Haddad, F. & Cooper, D. M. Impact of brief exercise on peripheral blood NK cell gene and microRNA expression in young adults. J Appl Physiol (1985) 114, 628–636 (2013). 10.1152/japplphysiol.01341.2012

66 Ramasamy, S., Velmurugan, G., Shanmugha Rajan, K., Ramprasath, T. & Kalpana, K. MiRNAs with apoptosis regulating potential are differentially expressed in chronic exercise-induced physiologically hypertrophied hearts. PLoS One 10, e0121401 (2015). 10.1371/journal.pone.0121401

67 Cengiz, M. et al. Differential expression of hypertension-associated microRNAs in the plasma of patients with white coat hypertension. Medicine (Baltimore) 94, e693 (2015). 10.1097/MD.0000000000000693

68 Ferland-McCollough, D. et al. Programming of adipose tissue miR-483-3p and GDF-3 expression by maternal diet in type 2 diabetes. Cell Death Differ 19, 1003–1012 (2012). 10.1038/cdd.2011.183

69 Mao, J., Li, C., Zhang, Y., Li, Y. & Zhao, Y. Human with-no-lysine kinase-4 3’-UTR acting as the enhancer and being targeted by miR-296. Int J Biochem Cell Biol 42, 1536–1543 (2010). 10.1016/j.biocel.2010.06.006

70 Qiao, Y. et al. miR-483-3p regulates hyperglycaemia-induced cardiomyocyte apoptosis in transgenic mice. Biochem Biophys Res Commun 477, 541–547 (2016). 10.1016/j.bbrc.2016.06.051

71 Torma, F. et al. The roles of microRNA in redox metabolism and exercise-mediated adaptation. J Sport Health Sci 9, 405–414 (2020). 10.1016/j.jshs.2020.03.004

72 Wang, T. et al. NFATc3-dependent expression of miR-153-3p promotes mitochondrial fragmentation in cardiac hypertrophy by impairing mitofusin-1 expression. Theranostics 10, 553–566 (2020). 10.7150/thno.37181

73 Yan, M. et al. miR-488-3p Protects Cardiomyocytes against Doxorubicin-Induced Cardiotoxicity by Inhibiting CyclinG1. Oxid Med Cell Longev 2022, 5184135 (2022). 10.1155/2022/5184135

74 Connolly, M. et al. miR-424-5p reduces ribosomal RNA and protein synthesis in muscle wasting. J Cachexia Sarcopenia Muscle 9, 400–416 (2018). 10.1002/jcsm.12266

75 de Gonzalo-Calvo, D. et al. Circulating microRNAs as emerging cardiac biomarkers responsive to acute exercise. Int J Cardiol 264, 130–136 (2018). 10.1016/j.ijcard.2018.02.092

76 de Gonzalo-Calvo, D. et al. Circulating inflammatory miRNA signature in response to different doses of aerobic exercise. J Appl Physiol (1985) 119, 124–134 (2015). 10.1152/japplphysiol.00077.2015

77 Fernandez-Sanjurjo, M. et al. Exercise dose affects the circulating microRNA profile in response to acute endurance exercise in male amateur runners. Scand J Med Sci Sports 30, 1896–1907 (2020). 10.1111/sms.13759

78 Rosa, A. et al. The interplay between the master transcription factor PU.1 and miR-424 regulates human monocyte/macrophage differentiation. Proc Natl Acad Sci U S A 104, 19849–19854 (2007). 10.1073/pnas.0706963104

79 Chai, F. et al. MicroRNA miR-181d-5p regulates the MAPK signaling pathway by targeting mitogen-activated protein kinase 8 (MAPK8) to improve lupus nephritis. Gene 893, 147961 (2024). 10.1016/j.gene.2023.147961

80 Nejad, C., Stunden, H. J. & Gantier, M. P. A guide to miRNAs in inflammation and innate immune responses. FEBS J 285, 3695–3716 (2018). 10.1111/febs.14482

81 Shah, R. et al. Small RNA-seq during acute maximal exercise reveal RNAs involved in vascular inflammation and cardiometabolic health: brief report. Am J Physiol Heart Circ Physiol 313, H1162–H1167 (2017). 10.1152/ajpheart.00500.2017

82 Zhang, Y. et al. MiR-181d-5p Targets KLF6 to Improve Ischemia/Reperfusion-Induced AKI Through Effects on Renal Function, Apoptosis, and Inflammation. Front Physiol 11, 510 (2020). 10.3389/fphys.2020.00510

83 Mononen, N. et al. Whole blood microRNA levels associate with glycemic status and correlate with target mRNAs in pathways important to type 2 diabetes. Sci Rep 9, 8887 (2019). 10.1038/s41598-019-43793-4

84 Arroyo, J. D. et al. Argonaute2 complexes carry a population of circulating microRNAs independent of vesicles in human plasma. Proc Natl Acad Sci U S A 108, 5003–5008 (2011). 10.1073/pnas.1019055108

85 Valadi, H. et al. Exosome-mediated transfer of mRNAs and microRNAs is a novel mechanism of genetic exchange between cells. Nat Cell Biol 9, 654–659 (2007). 10.1038/ncb1596

86 Vickers, K. C., Palmisano, B. T., Shoucri, B. M., Shamburek, R. D. & Remaley, A. T. MicroRNAs are transported in plasma and delivered to recipient cells by high-density lipoproteins. Nat Cell Biol 13, 423–433 (2011). 10.1038/ncb2210

87 Bosson, A. D., Zamudio, J. R. & Sharp, P. A. Endogenous miRNA and target concentrations determine susceptibility to potential ceRNA competition. Mol Cell 56, 347–359 (2014). 10.1016/j.molcel.2014.09.018

88 Mukherji, S. et al. MicroRNAs can generate thresholds in target gene expression. Nat Genet 43, 854–859 (2011). 10.1038/ng.905

89 Grimson, A. et al. MicroRNA targeting specificity in mammals: determinants beyond seed pairing. Mol Cell 27, 91–105 (2007). 10.1016/j.molcel.2007.06.017

90 Lim, L. P. et al. Microarray analysis shows that some microRNAs downregulate large numbers of target mRNAs. Nature 433, 769–773 (2005). 10.1038/nature03315

91 Peter, M. E. Targeting of mRNAs by multiple miRNAs: the next step. Oncogene 29, 2161–2164 (2010). 10.1038/onc.2010.59

92 Baek, D. et al. The impact of microRNAs on protein output. Nature 455, 64–71 (2008). 10.1038/nature07242

93 Selbach, M. et al. Widespread changes in protein synthesis induced by microRNAs. Nature 455, 58–63 (2008). 10.1038/nature07228

94 Krustrup, P., Ferguson, R. A., Kjaer, M. & Bangsbo, J. ATP and heat production in human skeletal muscle during dynamic exercise: higher efficiency of anaerobic than aerobic ATP resynthesis. J Physiol 549, 255–269 (2003). 10.1113/jphysiol.2002.035089

95 Smith, J. A. B., Murach, K. A., Dyar, K. A. & Zierath, J. R. Exercise metabolism and adaptation in skeletal muscle. Nature Reviews Molecular Cell Biology 24, 607–632 (2023). 10.1038/s41580-023-00606-x

96 Zagatto, A. M. et al. Impacts of high-intensity exercise on the metabolomics profile of human skeletal muscle tissue. Scand J Med Sci Sports 32, 402–413 (2022). 10.1111/sms.14086

97 He, A., Zhu, L., Gupta, N., Chang, Y. & Fang, F. Overexpression of micro ribonucleic acid 29, highly up-regulated in diabetic rats, leads to insulin resistance in 3T3-L1 adipocytes. Mol Endocrinol 21, 2785–2794 (2007). 10.1210/me.2007-0167

98 Hung, Y. H. et al. Acute suppression of insulin resistance-associated hepatic miR-29 in vivo improves glycemic control in adult mice. Physiol Genomics 51, 379–389 (2019). 10.1152/physiolgenomics.00037.2019

99 Massart, J. et al. Altered miR-29 Expression in Type 2 Diabetes Influences Glucose and Lipid Metabolism in Skeletal Muscle. Diabetes 66, 1807–1818 (2017). 10.2337/db17-0141

100 Pinto-Hernandez, P. et al. Training-induced plasma miR-29a-3p is secreted by skeletal muscle and contributes to metabolic adaptations to resistance exercise in mice. Mol Metab 98, 102173 (2025). 10.1016/j.molmet.2025.102173

101 Iacomino, G. et al. The association of circulating miR-191 and miR-375 expression levels with markers of insulin resistance in overweight children: an exploratory analysis of the I.Family Study. Genes Nutr 16, 10 (2021). 10.1186/s12263-021-00689-1

102 Li, W. et al. MicroRNA-191 blocking the translocation of GLUT4 is involved in arsenite-induced hepatic insulin resistance through inhibiting the IRS1/AKT pathway. Ecotoxicology and Environmental Safety 215, 112130 (2021). 10.1016/j.ecoenv.2021.112130

103 de Klerk, J. A. et al. Circulating small non-coding RNAs are associated with the insulin-resistant and obesity-related type 2 diabetes clusters. Diabetes Obes Metab 26, 4375–4385 (2024). 10.1111/dom.15786

104 Satake, E. et al. Circulating miRNA Profiles Associated With Hyperglycemia in Patients With Type 1 Diabetes. Diabetes 67, 1013–1023 (2018). 10.2337/db17-1207

105 Patra, D. et al. Adipose tissue macrophage-derived microRNA-210-3p disrupts systemic insulin sensitivity by silencing GLUT4 in obesity. J Biol Chem 300, 107328 (2024). 10.1016/j.jbc.2024.107328

106 Patra, D. et al. miR-210-3p Promotes Obesity-Induced Adipose Tissue Inflammation and Insulin Resistance by Targeting SOCS1-Mediated NF-κB Pathway. Diabetes 72, 375–388 (2023). 10.2337/db22-0284

107 Dube, J. J., Allison, K. F., Rousson, V., Goodpaster, B. H. & Amati, F. Exercise dose and insulin sensitivity: relevance for diabetes prevention. Med Sci Sports Exerc 44, 793–799 (2012). 10.1249/MSS.0b013e31823f679f

108 Holloszy, J. O. Exercise-induced increase in muscle insulin sensitivity. J Appl Physiol (1985) 99, 338–343 (2005). 10.1152/japplphysiol.00123.2005

109 Xu, G. et al. MiR-26b modulates insulin sensitivity in adipocytes by interrupting the PTEN/PI3K/AKT pathway. Int J Obes (Lond) 39, 1523–1530 (2015). 10.1038/ijo.2015.95

110 Houshmand-Oeregaard, A. et al. Increased expression of microRNA-15a and microRNA-15b in skeletal muscle from adult offspring of women with diabetes in pregnancy. Hum Mol Genet 27, 1763–1771 (2018). 10.1093/hmg/ddy085

111 Tao, H. et al. MiR-126 Suppresses the Glucose-Stimulated Proliferation via IRS-2 in INS-1 beta Cells. PLoS One 11, e0149954 (2016). 10.1371/journal.pone.0149954

112 Zhou, T. et al. Regulation of Insulin Resistance by Multiple MiRNAs via Targeting the GLUT4 Signalling Pathway. Cell Physiol Biochem 38, 2063–2078 (2016). 10.1159/000445565

113 Aravin, A. A. et al. A piRNA pathway primed by individual transposons is linked to de novo DNA methylation in mice. Mol Cell 31, 785–799 (2008). 10.1016/j.molcel.2008.09.003

114 Girard, A., Sachidanandam, R., Hannon, G. J. & Carmell, M. A. A germline-specific class of small RNAs binds mammalian Piwi proteins. Nature 442, 199–202 (2006). 10.1038/nature04917

115 Iwasaki, Y. W., Siomi, M. C. & Siomi, H. PIWI-Interacting RNA: Its Biogenesis and Functions. Annu Rev Biochem 84, 405–433 (2015). 10.1146/annurev-biochem-060614-034258

116 Weick, E. M. & Miska, E. A. piRNAs: from biogenesis to function. Development 141, 3458–3471 (2014). 10.1242/dev.094037

117 Mei, Y. et al. A piRNA-like small RNA interacts with and modulates p-ERM proteins in human somatic cells. Nat Commun 6, 7316 (2015). 10.1038/ncomms8316

118 Ozata, D. M., Gainetdinov, I., Zoch, A., O’Carroll, D. & Zamore, P. D. PIWI-interacting RNAs: small RNAs with big functions. Nat Rev Genet 20, 89–108 (2019). 10.1038/s41576-018-0073-3

119 Zhang, D. et al. The piRNA targeting rules and the resistance to piRNA silencing in endogenous genes. Science 359, 587–592 (2018). 10.1126/science.aao2840

120 Ingerslev, L. R. et al. Endurance training remodels sperm-borne small RNA expression and methylation at neurological gene hotspots. Clin Epigenetics 10, 12 (2018). 10.1186/s13148-018-0446-7

121 Harrell, J. F. E. in Springer Series in Statistics, 1 online resource (XXV, 582 pages 157 illustrations, 553 illustrations in color (Springer International Publishing : Imprint: Springer,, Cham, 2015).

122 Vittinghoff, E. Regression methods in biostatistics : linear, logistic, survival, and repeated measures models. 2nd edn, (Springer, 2012).

123 Loganantharaj, R. & Randall, T. A. The Limitations of Existing Approaches in Improving MicroRNA Target Prediction Accuracy. Methods Mol Biol 1617, 133–158 (2017). 10.1007/978-1-4939-7046-9_10

124 Alles, J. et al. An estimate of the total number of true human miRNAs. Nucleic Acids Res 47, 3353–3364 (2019). 10.1093/nar/gkz097

125 Diener, C., Keller, A. & Meese, E. The miRNA-target interactions: An underestimated intricacy. Nucleic Acids Res 52, 1544–1557 (2024). 10.1093/nar/gkad1142

